# Deep Texture Representations as a Universal Encoder for Pan-cancer Histology

**DOI:** 10.1101/2020.07.28.224253

**Authors:** Daisuke Komura, Akihiro Kawabe, Keisuke Fukuta, Kyohei Sano, Toshikazu Umezaki, Hirotomo Koda, Ryohei Suzuki, Ken Tominaga, Hiroki Konishi, Shu Nishida, Genta Furuya, Hiroto Katoh, Tetsuo Ushiku, Masahi Fukayama, Shumpei Ishikawa

**Author notes:** Lead Contact Further information and requests for resources should be directed to and will be fulfilled by the Lead Contact, Shumpei Ishikawa.

## Abstract

Cancer histological images contain rich biological and clinical information, but quantitative representation can be problematic and has prevented direct comparison and accumulation of large-scale datasets. Here we show that deep texture representations (DTRs) produced by a bilinear Convolutional Neural Network, express cancer morphology well in an unsupervised manner, and work as a universal encoder for cancer histology. DTRs are useful for content-based image retrieval, enabling quick retrieval of histologically similar images from optimally area selected datasets of 7,175 cases from The Cancer Genome Atlas. Via comprehensive comparison with driver and clinically actionable gene mutations, we have successfully predicted 309 combinations of genomic features and cancer types from hematoxylin and eosin-stained images at high accuracy (AUC > 0.70 and q < 0.02). With its mounting capabilities on accessible devices such as smartphones, DTR-based encoding for cancer histology has a potentially strong impact on global equalization for cancer diagnosis and targeted therapies.

## Introduction

For more than a century, pathologists have observed tissue slides via microscope not only to diagnose diseases but also to uncover clues for understanding disease pathophysiology. Careful observation has led to the discovery of numerous biologically and clinically relevant morphological features of both pathogens and host tissues. However, one critical limitation of this approach is that there has been no way to objectively and quantitatively express histological features, making it difficult to communicate with others among research/clinical communities, and to compare the data directly with other cases. This situation strongly contrasts with that of genomic data, for which researchers can directly compare as well as accumulate and retrieve new findings from thousands of cases. As such, investigations like genome-wide association studies (GWAS) have a substantial impact on current biological and clinical sciences.

Application of the Convolutional Neural Network (CNN), a deep-learning technology, has thus far been successful for classification, detection, and segmentation of histopathology images. CNN has already surpassed human pathologists in terms of accuracy and diagnostic time within some applications (Bejnordi et al., 2017; Nagpal et al., 2018). However, even though this approach is powerful, CNNs require a task-specific feature extraction process for individual tasks, meaning that a huge amount of training data is essential to construct trained networks to cover every subject in histopathology fields. This setback hampers the widespread application of artificial intelligence in pathology communities. Another way to utilize CNN within histopathology is via use as a general feature extractor (Hatipoglu and Bilgin, 2016). However, unlike general images like those of cats and dogs, morphological features of histopathology images—especially for cancer histology—consist of spatial invariance, such as the texture(s) derived from cellular and extracellular components, rather than their relative positions within the image as is observed in typical photographs, for example (Masaki, 2006).

This phenomenon has prompted us to develop an unsupervised feature extractor which can express tissue morphology objectively and quantitatively (Figure 1). We show that DTRs produced by a bilinear CNN (B-CNN) (Lin et al., 2015) could adequately express cancer morphology information in an unsupervised manner and function as a universal encoder for cancer histology. DTRs are based on a B-CNN pre-trained with general images for feature extraction (Figures 1A and 1B), which enables extraction of spatially invariant features independent of the relative location of individual cellular and extracellular components within the image. We then used compact bilinear pooling (CBP) (Gao et al., 2015) to reduce the dimensions of the image representation, which enables quick processing in practical settings. DTRs were applied to 8,736 hematoxylin and eosin (H&E)-stained histopathology whole slide images (WSIs) of 32 cancer types from The Cancer Genome Atlas (TCGA) (Weinstein et al., 2013) (Table S1), and validated these within two clinically important applications: 1) content-based image retrieval (CBIR), i.e. “identification of histologically similar images among large datasets,” and 2) prediction of genomic features (e.g., driver gene mutations) from histology images. After selection of uniform cancerous regions by trained pathologists, 271,700 patches with 128 × 128 µm were extracted and resized to 256 × 256 pixels for further analysis.

**Figure 1.**
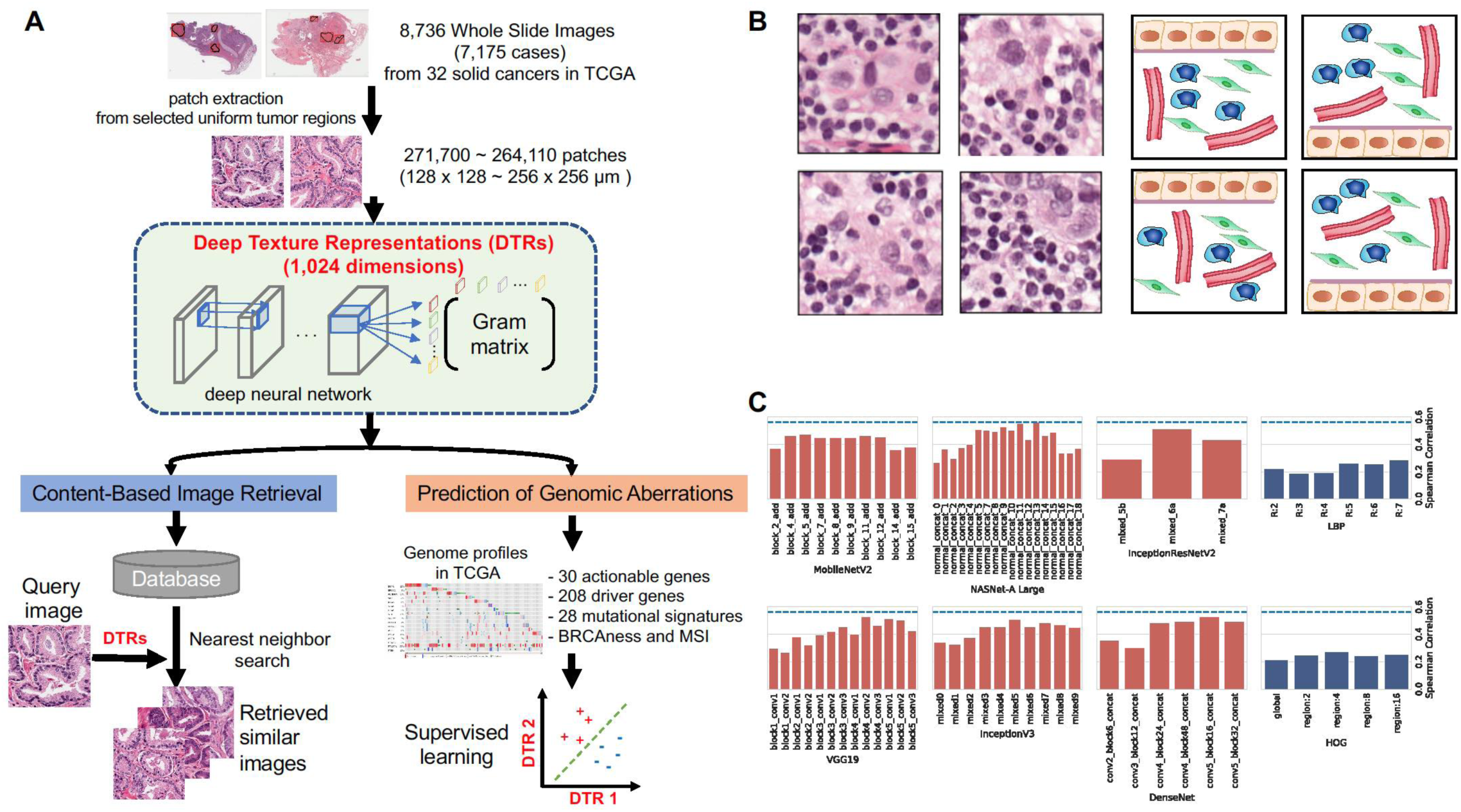
Unsupervised encoding of cancer histology by deep texture representation (DTRs). (A) Workflow overview. Uniform tumor regions within 8,736 whole slide images representing 32 solid cancers from TCGA dataset were manually annotated by trained pathologists, and 256 × 256 pixel patches were generated. By utilizing the optimal pre-trained CNN, 1,024-dimensional vector representations were extracted for each patch and tested in two applications: CBIR to retrieve histologically similar images, and prediction of genomic aberration from histopathology images. (B) Four histologically similar representative images (left) and pattern diagrams (right) with positional variance but similar deep texture. The extracted features are invariant to the relative positions of the components in the images, which is essential for histopathological image analysis. (C) Spearman’s rank correlation between similarity of images ranked by pathologists and DTRs. The layers are deeper along the x-axis from left to right in each network. Middle layers in NasNet-A Large network showed the strongest association to similarity ranked by pathologists.

## RESULTS

### Finding an optimized deep neural network for DTR of cancer histology

The architecture of the neural network and the layers used within bilinear operations affects extracted features. Thus, we first determined the optimal combination of the network and layer for DTRs—which properly captures characteristics of histopathology images—by evaluating the correlations between similarity rankings of the images calculated using features obtained by DTRs from various representative pre-trained models, and the similarities ranked by trained pathologists (Figure 1C). The DTRs correlated well with pathologist analyses, and they outperformed conventional image features such as histogram of oriented gradient (HOG) (Dalal and Triggs, 2005) and local binary pattern (LBPs) (Ojala et al., 2002). Interestingly, the middle layers in most CNN architectures outperformed the first and last layers within the same network. It is suggested that as networks are trained using general images, and convolutional layers learn spatial pattern hierarchies, middle layers that capture intermediate-level features are better suited to histopathology image analysis as compared to first layers that learn low-level features (e.g., edges) in smaller receptive fields, or last layers that learn high-level features in larger receptive fields (Yosinski et al., 2015). We also compared DTR performance using 1,024, 8,192 and 32,768 dimensions for final representations, and as a result all DTRs exhibited comparable performance (Figure S1). Thus, the most compact (1,024 dimensions) was adopted as the final representation for each image in subsequent analyses in order to reduce computational time.

### DTRs of pan-cancer histology

When 7,175 TCGA pan-cancer cases are projected in two-dimensional space using their DTRs, we found that they are well distributed based on morphologic features, and that histologically similar images came in close proximity (Figure 2A). While images from different tissues occupied different areas in general, those of different tissue origin with similar histopathological appearances (e.g. squamous cell carcinomas from multiple organs) tended to overlap significantly within each category (Figures 2B and S2A). Importantly, patches with similar morphology but differing H&E stain intensity also tended to fall close to each other, indicating that DTR encoding is robust despite color variation, which is an issue that frequently poses a major problem when comparing cases from different hospitals (Figure S2B). In a detailed analysis, DTRs clearly showed differential distributions according to histological subtypes within each cancer category, as in the cases of pancreatic ductal/neuroendocrine adenocarcinoma, stomach intestinal/diffuse adenocarcinoma, and Gleason grade in prostate adenocarcinoma (Figure 2C).

**Figure 2.**
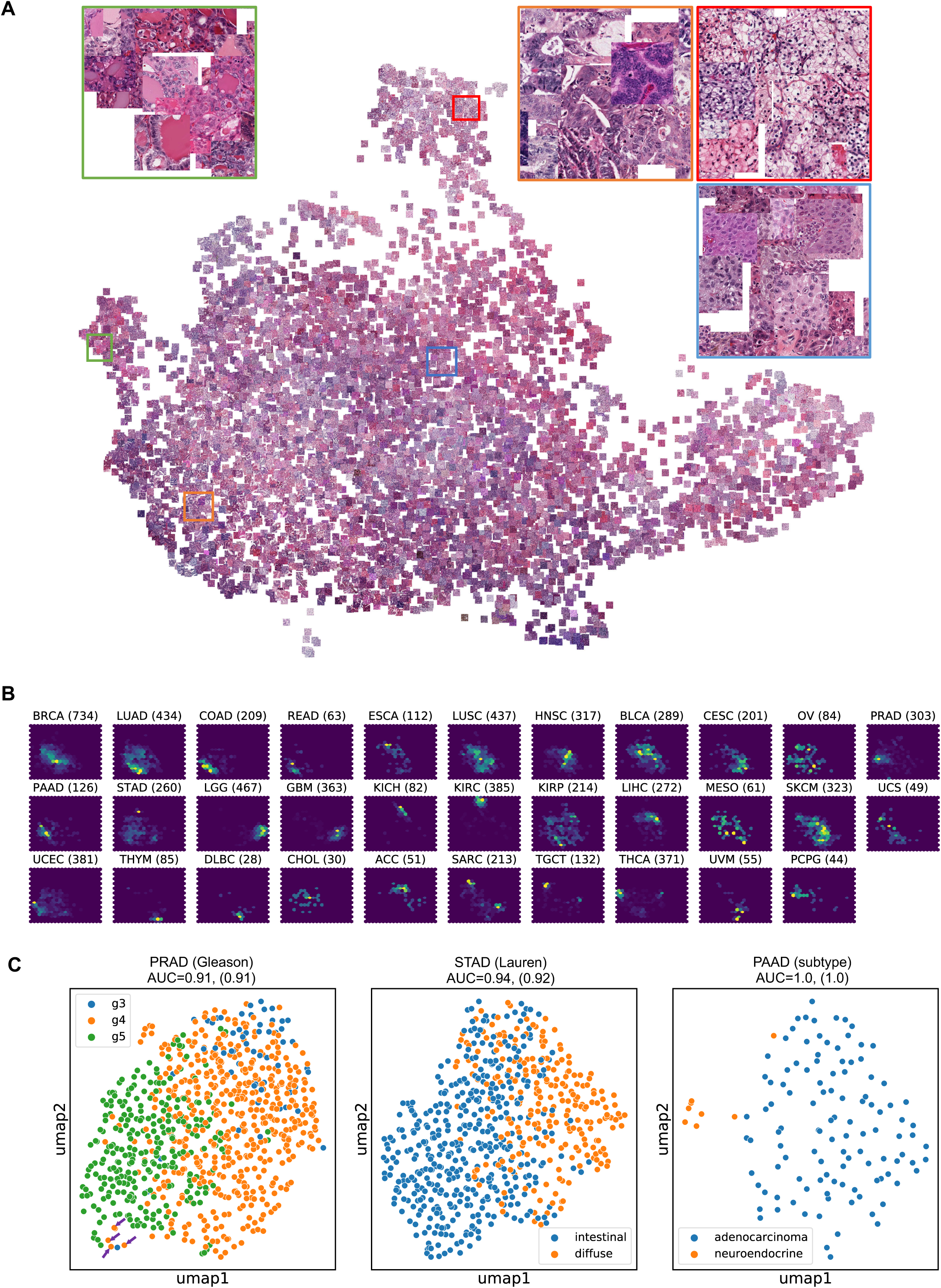
DTRs of pan-cancer histology. (A) 2D projection of the DTRs of TCGA pan-cancer histology images. The representative 7,175 images (128 × 128 µm) are shown at relative positions according to their DTR vectors using uniform manifold approximation and projection (UMAP). Enlarged images of the regions indicated by colored boxes are shown around the projection. (B) Distribution of each cancer type in UMAP projection. The numbers in parentheses following cancer type indicate the number of patients included. LUAD, lung adenocarcinoma; COAD, colon adenocarcinoma; READ, rectum adenocarcinoma; PAAD, pancreas adenocarcinoma; LUSC, lung squamous cell carcinoma; HNSC, head and neck squamous cell carcinoma; CESC, cervical squamous cell carcinoma (see Abbreviations for the full list). (C) UMAP projection of Gleason scores in prostate adenocarcinoma (left), Lauren classifications in stomach adenocarcinoma (middle), subtypes of pancreatic adenocarcinoma patients (right) for each region selected by the pathologists. Mean area under the curve (AUC) values based on logistic regression using DTRs without or with (indicated in parentheses) color normalization are also shown. Arrows within the Gleason score plot indicate a case with a Gleason score of 4 at outlier-like positions, which belong to a hypernephromatoid pattern subtype (Figure S2C). Color normalization did not improve classification performance, indicating the robustness of DTRs to color variation.

### Content-based image retrieval (CBIR) of histologically similar images by DTR

There has long been a need to utilize CBIR to retrieve histologically similar images for educational, diagnostic, and research purposes, and this technology could contribute significantly to the standardization of diagnostic quality. We attempted CBIR by retrieving the most relevant images from TCGA pan-cancer datasets based on the cosine similarity of DTR spaces and approximate nearest neighbor search method (Figure 1A). Using representative query images from multiple cancer categories, we found that the retrieved appeared histologically similar to the query images (Figure 3A). DTRs contributed to successful image retrieval for each query image from the same categories of 32 cancer types as defined by TCGA (precision: 90.8 ± 10% among the top 10 ranked images, and 78.8 ± 15.4% among the top 20 ranked images) (Figure 3B). Subsequently, we evaluated the concordance between the cosine similarity defined by DTRs and histopathological similarity judged by 5 independent trained pathologists in a strictly blinded comparison. As a result, images that the CBIR determined to be most similar to the query images and those selected by five pathologists were almost identical (59/60 tests), except for one case where the image judged to be the second most similar by the DTR was selected by one of the pathologists (Figure S3). These results demonstrated that DTRs adequately capture the morphologic features and function as a universal encoder for pan-cancer histology.

**Figure 3.**
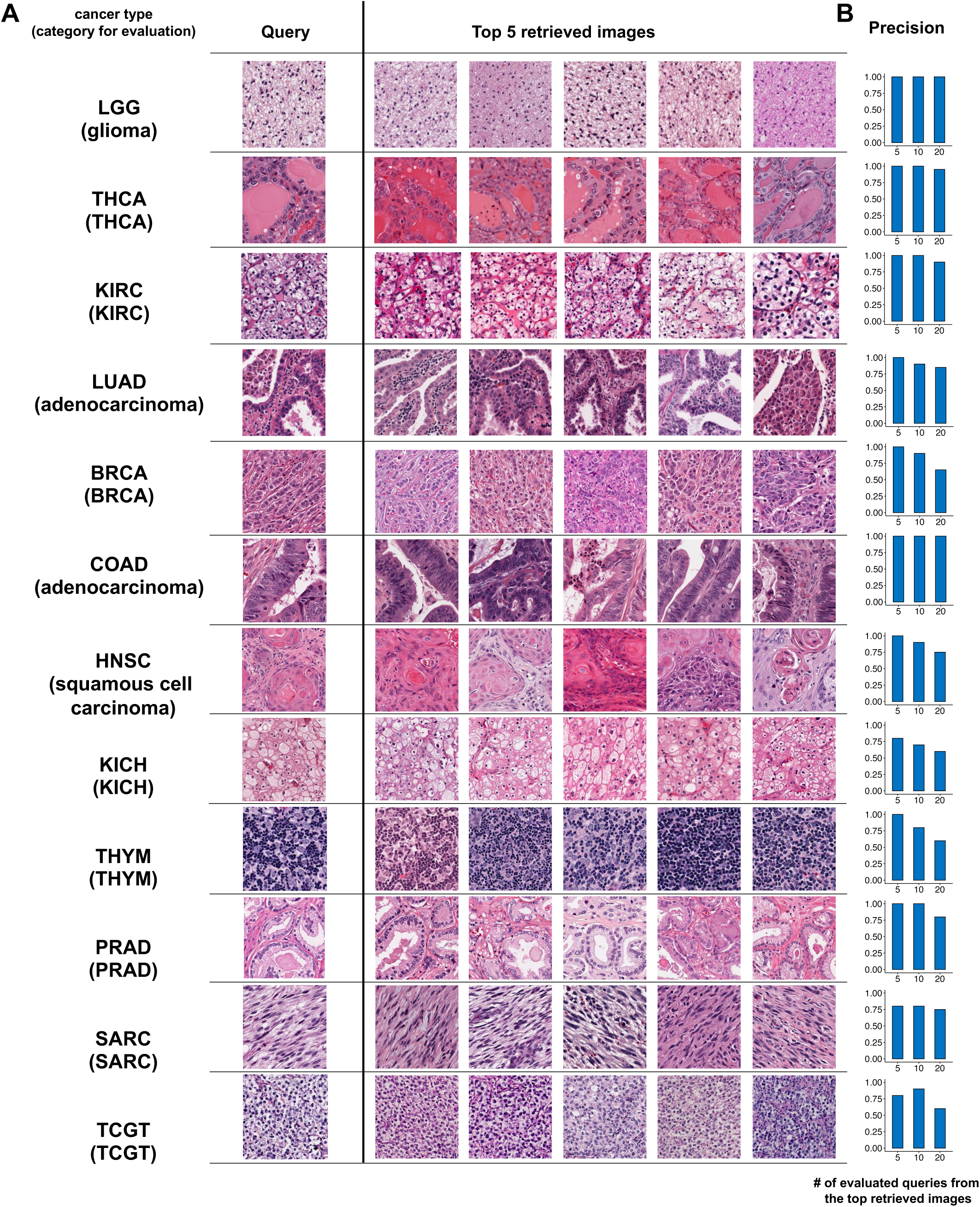
Content-based image retrieval of histologically similar images by DTR. (A) CBIR extraction of histologically similar images. Representative query images and top 5 retrieved results are shown for twelve cancer types. The five images were retrieved from the cancer histology images of 7,175 patients across all the cancer types in TCGA dataset based on the cosine similarity by DTRs. (B) Precision (number of images in the correct category / number of retrieved images) of CBIR for the 12 query images. Precision values based on the categories for top 5, 10 and 20 retrieved histopathology images are shown. The cancer type was used as the category except for LUAD, COAD, and HNSC, in which adenocarcinoma or squamous cell carcinoma were used as the ground truth category.

### Comprehensive evaluation of cancer genome-histology correlations by the DTRs of H&E image

Once DTRs are generated in an unsupervised manner, they can be correlated to other biological and clinical parameters. We therefore examined the correlation between histology and cancer genome aberrations, aiming to discover potential morphologic features induced by somatic cancer driver mutations. The correlation between DTRs and 208 informative somatic mutations—including single nucleotide variations (SNVs), indels, copy number variations (CNVs), and fusions—out of 322 cancer driver genes were investigated in OncoKB database (Chakravarty et al., 2017) using 1,024-dimensional DTRs as features for a logistic regression model. Stratified 3-fold cross-validation was performed, and a subset of these data were validated in our independent datasets outside of TCGA (see Supplementary Methods). As a result, we discovered that 309 genomic feature cancer type combinations, including 42 clinically actionable mutation status, exhibit strong correlation with DTRs and are predictable based solely on H&E histopathology images with high accuracy (AUC of receiver operating characteristics > 0.70, q < 0.02) (Figures 4A, 4B and S4A, Tables S2-S5). These include previously reported correlations such as *STK11* and *TP53* in lung adenocarcinoma (LUAD) (Coudray et al., 2018), *SPOP* in prostate adenocarcinoma (PRAD) (Schaumberg et al., 2017), as well as these coupled with known subclasses, such as *CDH1* mutations linked to lobular and diffuse-type gastric adenocarcinomas (Pharoah et al., 2001; Bass et al., 2014). However, to the best of our knowledge, no reports of a comprehensive evaluation of cancer classes and drivers have been made, and most of the correlations found within the current study are either novel or not previously described in detail. Uterine corpus endometrial carcinoma (UCEC) and stomach adenocarcinoma (STAD) contain morphologic correlations with many driver genes, thereby reflecting their major distinct biological subtypes (i.e. endometrioid/serous microsatellite instability (MSI) in UCEC and diffuse, intestinal, MSI or EBV^+^ in STAD). While some driver gene mutations are linked to the morphology of a specific cancer type—such as *GTF2* to– thymoma (THYM) and *NF2* to kidney renal papillary cell carcinoma (KIRP)—general cancer driver genes such as *TP53, CDKN2A/CDKN2B, PIK3CA, PTEN*, and *MYC*, which are responsible for well-known cancer hallmarks like genomic stability, cell cycle, and cellular metabolism (Hanahan and Weinberg, 2011), are linked to histologic features in a broad range of cancer types (Figure 4A and 4C). The performance of models trained on TCGA cases for the prediction of driver mutations is validated by our independent STAD datasets (Figures S4B–S4D).

**Figure 4.**
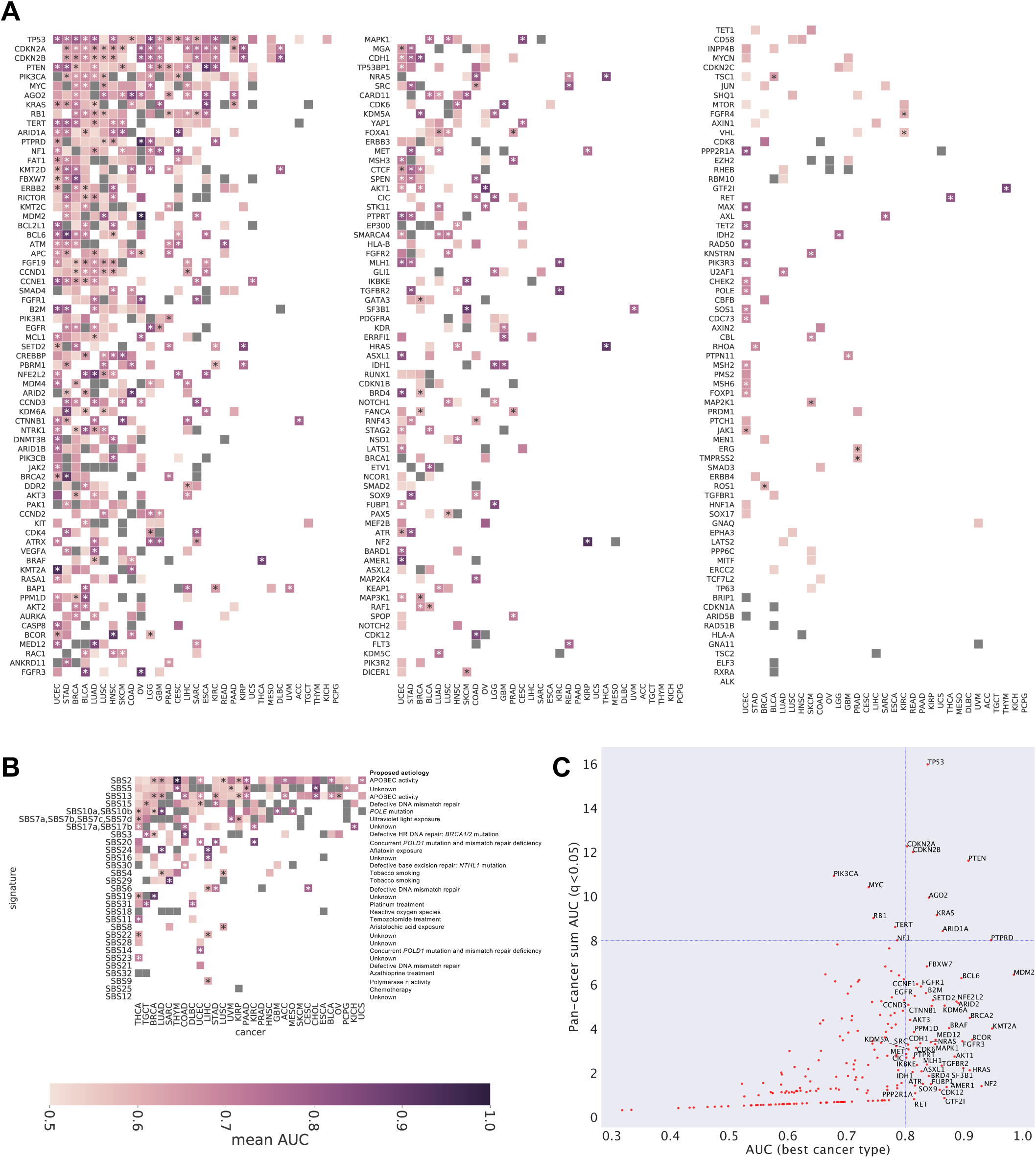
Prediction of cancer-related genomic mutations from the DTRs of H&E histology image. Area under the receiver operating characteristic (ROC) curves values for the prediction of somatic mutations of (A) OncoKB-annotated 208 driver genes, and (B) single base substitution (SBS) mutational signatures in 32 cancer types. Somatic SNVs, indels, CNVs and fusions are integrated in that their direction of gain- or loss-of-function does not contradict each other. Asterisks indicate q < 0.02; grey boxes indicate AUC < 0.5; white space indicates that the mutations could not be predicted due to a lack of adequate sample number for genomic aberrations. (C) A global view of pan-cancer and cancer type-specific correlations between morphology and genomic aberrations. X and y axes represent AUC values and total AUC values with q < 0.05 in the 32 cancer types, respectively.

### Correlation of morphological features with somatic genomic mutations

We then investigated what morphological changes are induced by these driver gene mutations. Comparison of top- and bottom-scored images for individual driver genes revealed distinct morphologic characteristics in mutant-positive cases (Figures 5A and S5A), which was even recognizable upon visual inspection by non-pathologists. Some of these features can also be expressed with classical histopathologic keywords (Table S6); for example, *FGFR3-*mutated bladder urothelial carcinoma (BLCA) exhibits regularly arranged cells with thin vascular stoma, and *CTNNB1*-mutated skin cutaneous melanoma (SKCM) exhibits sheet-like spindle cell tumors with clear melanin pigment production. By examining morphological changes induced by broad effectors in multiple cancer types, we have found that some oncogenic drivers induce universal morphologic change across cancer types. For example, the *TP53* mutation—which causes chromosomal instability—induces an increase in nuclear size and atypia within LUAD, breast invasive carcinoma (BRCA), UCEC, HNSC and glioblastoma multiforme (GBM) (Figure S5B). In contrast, *MYC* amplification induces an increase in cell size with plump cytoplasm, typically in liver hepatocellular carcinoma (LIHC), PAAD, LUSC, BRCA, and BLCA (Figure 5B). As MYC is a major accelerator of glycolytic metabolism, our result indicates a relationship between cell size regulation and cellular metabolism, which has also been suggested by several lines of experimental evidence (Baena et al., 2005; Zanet et al., 2005).

**Figure 5.**
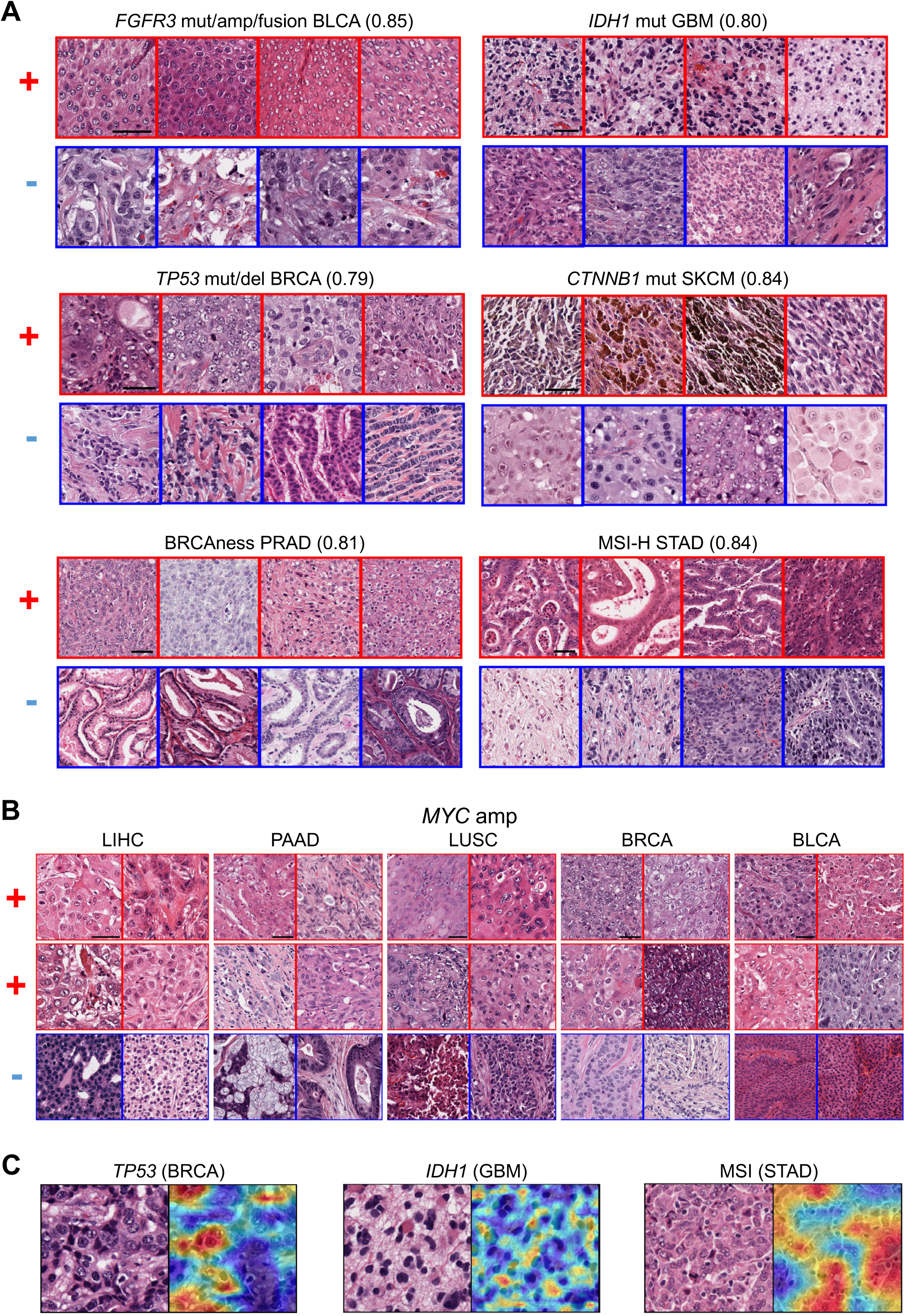
Morphologic features correlated with somatic genomic mutations. (A) Representative images of cases with high (top row) and low (bottom row) somatic mutation probability based on logistic regression models are shown for *FGFR3* in BLCA, *IDH1* in GBM, *TP53* in BRCA, *CTNNB1* in SKCM, BRCAness signature in PRAD, and MSI status in STAD. Images with high and low probabilities were selected from mutation-positive and -negative cases, respectively. The frame color indicates presence (red) or absence (blue) of mutations. All eight images shown for each gene category are of the same magnification. Numbers in parentheses indicate mean AUC of the prediction for each cancer-mutation combination in 3-fold cross-validation. Scale bar: 50 µm. (B) Morphological changes accompanying *MYC* amplification in different cancer types. Increased cell size with pump cytoplasm is observed for LIHC, PAAD, LUSC, BRCA, and BLCA. All images are shown in the same magnification. Scale bar: 50 µm. (C) Grad-CAM visualization of mutation-related morphological changes by CNN. *TP53* and *IDH1* mutations are primarily linked with the nucleus and cytoplasm of cancer cells, respectively, and MSI status is overlapped with tumor-infiltrating lymphocytes.

In some cancer type-driver gene combinations with large sample sizes and appropriate class balance to be fed into the learning of deep neural network, we successfully obtained an objective depiction of the biologically reasonable, mutant-specific morphologic features by Grad-CAM approach (Figure 5C, see Supplementary Methods). While *TP53* mutations in breast cancer are linked to atypical cancer nuclei, *IDH1* mutations in GBM are linked not to nuclear features but instead to astrocyte-like faint or gemistocytic cytoplasmic features, reflecting that *IDH1*-mutant GBM developed from low-grade astrocytomas (Cancer Genome Atlas Research Network et al., 2015). Mutation signatures associated with MSI in STAD overlap with intraepithelial lymphocytic infiltration, reflecting the well-established link between neoantigens and intratumor lymphocytic infiltration (Phillips et al., 2004). Within thyroid carcinoma (THCA), we observed a clear link between oncogenic pathway mutation and specific morphology; *BRAF*, RASs *(HRAS, NRAS)*, and RTKs (*RET*, NTRKs, *ALK*, etc.) mutant groups exhibited group-specific morphologic features, indicating that similar pathway signals generate similar tumor morphology and biology (Figure 6A-6C).

**Figure 6.**
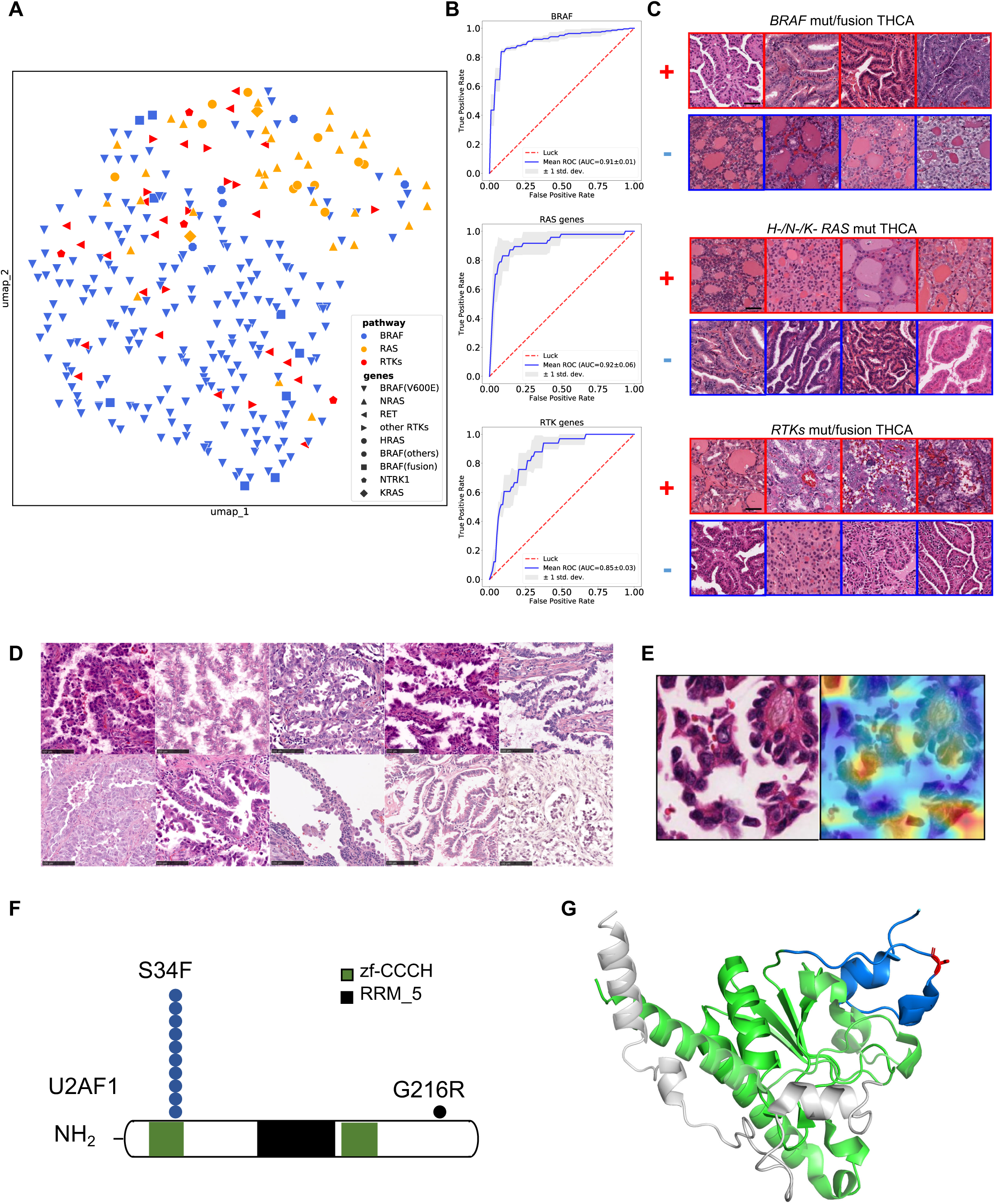
Pathway-level morphological correlation and a biologically distinct subclass, discovered by DTR-based comprehensive evaluation of cancer histology. (A) Distribution of histology images of THCA cases within UMAP projection. Three pathway mutations are differentially as indicated by the colored dots: *BRAF, H-/N-/K-RAS*, and receptor tyrosine kinases (RTKs) (*RET, NTRK*s, and *ALK*). Dot shape indicates the mutated gene in each case. *BRAF*- and *X-RAS*-mutated cases are differentially positioned, and many RTK-mutated cases fall in the intermediate zone. (B) The cross-validation performance of the logistic regression models using DTRs was measured in terms of AUC for *BRAF, RAS (N-RAS/H-RAS/K-RAS)*, and *RTK* mutations. AUC values in *RAS*s and RTKs were larger when measured altogether versus when measured individually (0.91 for *H-RAS* and 0.89 for *N-RAS*; 0.77 for *RET* and 0.76 for *NTRK1*). (C) Papillary structure with tall cells is observed in *BRAF*-mutated cases, while H-RAS- and N-RAS-mutated cases showed follicular morphology. RTK-mutated cases exhibit structures in which the epithelial front and back are reversed. All images for each gene-cancer combination are shown with the same magnification. Both somatic SNV and fusion were taken into account in (B) and (C). Scale bar: 50 µm. (D-F) A novel distinct lung adenocarcinoma subclass with splice-complex protein *U2AF1* mutations. (D) *U2AF1-*mutated cases share similar histology of differentiated lepidic predominant adenocarcinoma with hobnail-type cells. Scale bar: 100 µm. (E) Grad-CAM visualization highlights the hobnail appearance of epithelial projections as a feature linked to *U2AF1* mutations. (F) All cases, with one exception, had an S34F hotspot mutation on the CCCH zinc finger domain, which strongly suggests its driver nature in this subgroup. (G) The 3D position of S34, which is highly conserved across species, is displayed using the crystal structure of the yeast U2AF complex (PDB: 4YH8). The red colored S34 of U2AF23 (yeast orthologue of human U2AF1) (green), is on the surface of conserved zf-CCCH domain (marine blue), distant from contact site with U2AF59 (yeast orthologue of human U2AF2) (grey).

Through these studies, we found distinct and previously undescribed subgroups with characteristic genomic and morphologic features. U2AF1 is a component of a spliceosomal complex, and LUAD with *U2AF1* mut is a well-differentiated adenocarcinoma exhibiting distinct lepidic predominant tumor with a hobnail appearance of the tumor cells (Figure 6D–6G). Almost all of the cases (10/11) had an S34F hotspot mutation, suggesting a gain-of-function nature for the mutant proteins. Serine at 34-th amino acid is highly conserved across species, which is on the surface of the conserved zf-CCCH domain and distant from contact site with another U2AF complex member. CTNNB1 is a transcription factor complex involved in adrenal development (Berthon et al., 2012) and *CTNNB1-*mutated adrenocortical carcinoma (ACC) comprises regularly packed cells with round nuclei (Figure S5C). All CTNNB1 mutations found in this subgroup were at mutation hot spot in exon3. Although pathologists can note with relative ease the correlation of morphology and specific genomic features once presented, evaluation of quantified cancer histology such as DTRs makes it practically possible to survey all possible combinations of mutations and cancer types comprehensively.

## DISCUSSION

CBIR and mutation prediction from routine H&E images have broad clinical implications. As most pathologists manually refer to a histology atlas when encountering unknown cases, CBIR would drastically decrease the time required for correct diagnosis. This is especially useful in small to medium-sized hospitals and those in developing countries, where pathologists often work alone without any practical access to consults. Our studies also uncovered 42 clinically actionable mutations and genomic features that are predictable with high accuracy (AUC > 0.70) from H&E images, including: *EGFR, ERBB2, FGFR1/2/3, PIK3CA, IDH1/2, BRAF, KIT*, and BRCAness as well as MSI status. Although this method is not 100% accurate and requires genetic testing for definitive diagnosis, systematic selection of highly probable candidates is important in many contexts. Particularly in developing countries where clinical resources are limited, it could lead to a substantial reduction of negative results, and resultant total number of subsequent costly cancer gene panel tests at the population level. To facilitate DTR-based analysis for routine H&E histologic images, we implemented our program both to a web server (Figure 7A) and smartphone devices (Figure 7B and 7C, Supplementary Video 1). By using a smartphone microscope device, clinicians not equipped with a digital slide camera device can take images and access our developed image analysis program. Effective dimensionality reduction of DTRs successfully enabled a “query-to-display time” of less than 4 seconds, which is reasonable in routine clinical settings.

**Figure 7.**
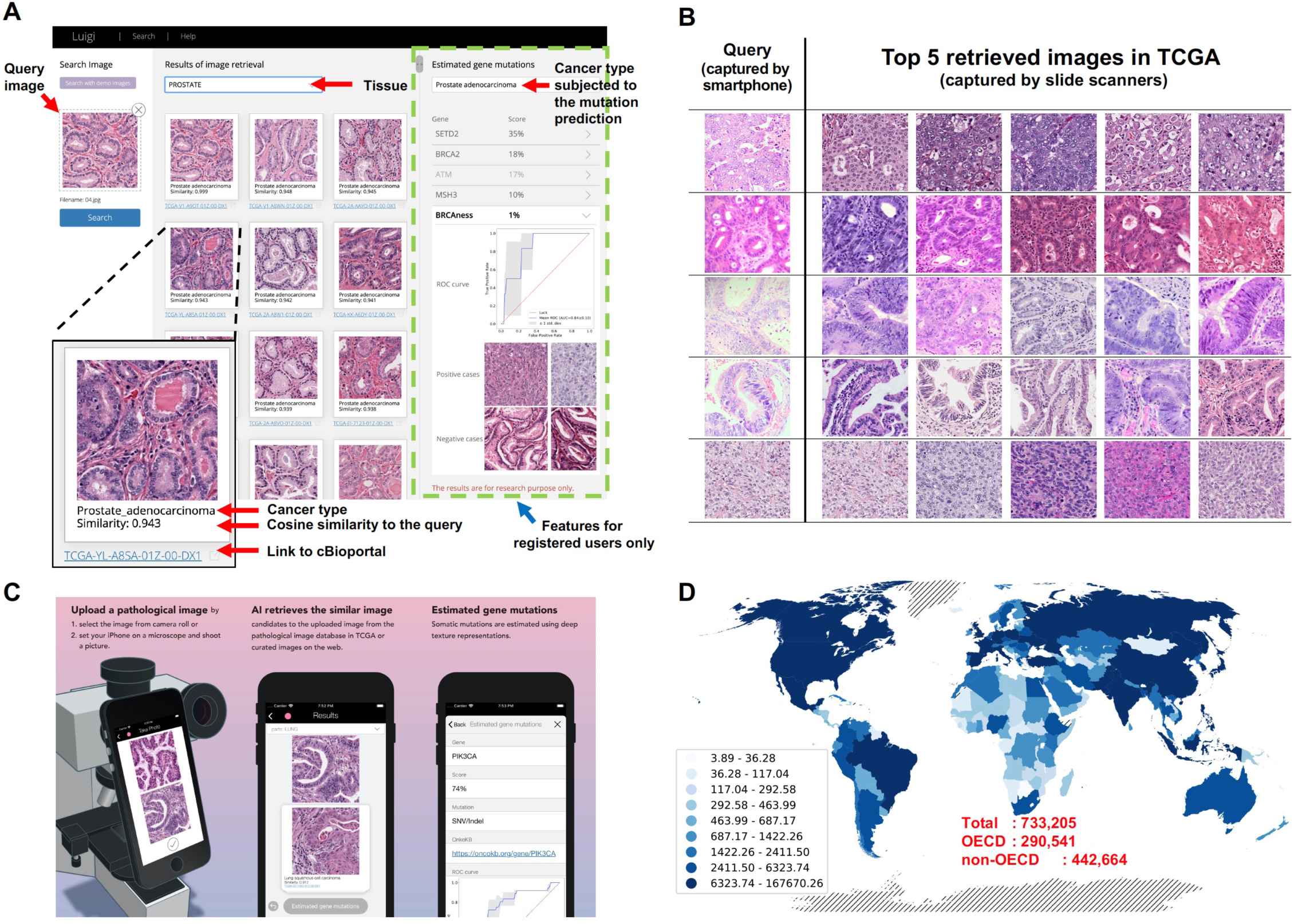
Global impact of CBIR and actionable mutation prediction with its implementation in an easily accessible environment. (A) Users upload a histopathological image either by dragging and dropping the file, pasting a screenshot of the image, or clicking “click here to select a file”. The Luigi system will retrieve the most relevant patch in each WSI, rank each patch according to cosine similarity, and initially display the relevant cancer cases among all cancer types. Tissues of origin can be changed by selecting the tissue from the pull-down menu above the results. By clicking the TCGA slide ID below each image, users can display the summary of the selected case on the cBioPortal website (https://www.cbioportal.org/), which includes the pathology report as well as clinical and molecular information for the case. The right panel shows the predicted genomic aberrations. (B) CBIR using query images acquired with a smartphone to extract histologically similar images. Examples of query images and the top 5 resulting retrieval results corresponding to each query image are shown. (C) User interface and usage of CBIR smartphone application. Users upload original cancer histology images captured using a smartphone camera, and histologically similar images and predicted mutations are displayed. (D) A global map showing the estimation of annual cancer cases with clinically actionable mutation which can be identified with our procedure with > 70% positive prediction value (see Supplementary Methods). The total number of cases found to be positive annually using our method in OECD and non-OECD countries are shown in red.

Our studies have unveiled the link between common actionable mutations and DTRs in major cancers such as: LUAD, BRCA, COAD, PRAD, and STAD. Our rough estimation by interpolating the actionable mutation frequency in TCGA to global cancer statistics indicates that approximately 0.73 million cancer cases with positive actionable mutations can be identified with a positive prediction value of > 70% each year, of which 0.44 million cases are from non-Organization for Economic Co-operation and Development (OECD) developing countries (Figures 7D and S6A, see Supplementary Methods). Although this is the theoretical maximum under the assumption that most cases have equal access to routine pathological evaluation, more than 40 cancer cases can be identified per one thousand patients within both OECD and non-OECD countries by the same criteria (Figure S6B). When the statistics are confined to cases treatable with small molecule drugs that can easily be stored and transported (e.g. BRCAness signatures correlated with platinum drug efficacy), 0.56 million cases worldwide (of which 0.35 million are from non-OECD countries) can be identified at the maximum (Figures S6C and S6D, Table S7).

The deep texture representations used in this study encompass several practical advantages: 1) they function without requiring extra learning with histopathology images, thus can be applied to relatively small datasets, such as rare disease categories, and 2) computational cost is relatively small once the features are extracted, making rapid decisions possible, as presented here. Although our study is limited to cancer histology, it is possible to extend this encoding method to the non-cancer fields as well. By acquiring a universal quantitative encoder for routinely-used H&E histology, pathologists can collect, accumulate, analyze, and query cases similar to systems for genomic or text information, which would have a profound impact on biomedicine and global health.

## Abbreviations

Abbreviations: Cancer type
ACC: Adrenocorticalcarcinoma
BLCA: Bladder_Urothelial_Carcinoma
LGG: Brain_Lower_Grade_Glioma
BRCA: Breast_invasive_carcinoma
CESC: Cervical_squamous_cell_carcinoma_and_endocervical_adenocarcinoma
CHOL: Cholangiocarcinoma
COAD: Colonadenocarcinoma
ESCA: Esophagealcarcinoma
GBM: Glioblastomamultiforme
HNSC: Head_and_Neck_squamous_cell_carcinoma
KICH: Kidney Chromophobe
K I RC: Kidney renal clear cell carcinoma
KIRP: Kidney_renal_papillary_cell_carcinoma
LIHC: Liver_hepatocellular_carcinoma
LUAD: Lungadenocarcinoma
LUSC: Lung_squamous_cell_carcinoma
DLBC: Lymphoid_Neoplasm_Diffuse_Large_B-cell_Lymphoma
MESO: Mesothelioma
OV: Ovarian_serous_cystadenocarcinoma
PAAD: Pancreaticadenocarcinoma
PCPG: Pheochromocytoma_and_Paraganglioma
PRAD: Prostateadenocarcinoma
READ: Rectumadenocarcinoma
SARC: Sarcoma
SKCM: Skin_Cutaneous_Melanoma
STAD: Stomachadenocarcinoma
TGCT: Testicular_Germ_Cell_Tumors
THYM: Thymoma
THCA: Thyroid carcinoma
UCS: UterineCarcinosarcoma
UCEC: Uterine_Corpus_Endometrial_Carcinoma
UVM: UvealMelanoma

## Methods

### TCGA whole slide image dataset

WSIs of TCGA dataset representing 32 solid cancer types were downloaded from Genomics Data Commons (GDC) legacy database from December 1, 2016 to June 19, 2017. In total, 9,662 diagnostic slides (with filename containing ‘DXn,’ where n represents the slide number) from 7,951 patients in SVS format were then processed for annotation. The number of slides for each cancer type are shown in Table S1.

### Image preprocessing

For each slide, at least three representative tumor regions were selected as polygons by two trained pathologists using a web browser-based software developed for this purpose. To select regions suitable for DTRs, uniform tumor regions were selected while avoiding as much as possible any regions with noncancerous structures. As a result, 926 slides were removed either due to poor staining, low resolution, being out of focus, lacking cancerous regions, or consisting of incorrect cancer types. As a result, 8,736 diagnostic slides from 7,175 patients were used for downstream analysis.

For DTR visualization, 10 patches from each of the annotated regions were arbitrarily cropped with random angles at 6 magnification levels (from 128 × 128 to 256 × 256 µm) using keras-OpenSlideGenerator (https://github.com/quolc/keras-OpenSlideGenerator). The pixel size corresponding to S µm was calculated as “, where *m* is micron per pixel (mpp), ranging from 0.11504 to 0.504 with a median value of 0.252. The mpp values were included as metadata (‘aperio.MPP’ or ‘openslide.mpp-x’) in the SVS files. Images with regions too small to crop images were skipped. Forty-two WSIs lacking mpp information were removed. Each patch was selected so as not to include areas outside the annotated region; selected regions were resized to 256 × 256 pixels. Consequently, the number of patches subjected to further analysis ranged from 264,110–271,700. For the CBIR application, the number of patches per region was limited to three to reduce memory consumption and search time for the system. As a result, 475,701 total patches were included in the database.

For the genome prediction tasks, the same dataset used within DTR visualization was utilized, but adenocarcinoma cases of CESC and ESCA were removed to avoid classification bias due to major histological category (e.g. adenocarcinoma and squamous cell carcinoma). The resulting number of patches ranged from 261,750–269,250.

### Deep texture representations (DTRs)

DTRs were computed by aggregating the averages of the location-wise outer products of the deep features in a CNN layer as follows:

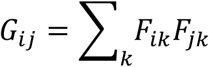

Where *G, F*, and *k* represent the Gram matrix, feature maps of the CNN, and position in the images, respectively. This manipulation—also known as bilinear pooling—produces orderless image representations. This process results in the discarding of some spatial information, and thus the representations contain spatial invariance, which appears to be suitable for cancer histology images as the similarity between two such images should not be affected by the relative location of cellular and extracellular components. In addition, as the representations are too high dimensional for practical application, CBP using Random Maclaurin approximation (Gao et al., 2015) was adopted to estimate the original high dimensional representations in lower final dimensions. CBP is an approximation bilinear pooling method utilizing two random matrices, and unlike other dimensionality reduction methods, such as principal component analysis, it does not require pre-training. Prior to the final representation, we added an element-wise signed square root layer and an instance-wise normalization as previously shown elsewhere (Lin et al., 2015).

### Selection of optimal neural network architecture and layer

DTRs can be computed on any convolutional layers in any CNN structure. Due to a lack of CNNs pre-trained with a large number of histopathology images, we have employed a model trained with an ImageNet dataset (Deng et al., 2009), which comprises over 10 million images with 1,000 classes and exhibits satisfactory performance within histopathology image analysis. To select an optimal network architecture and a corresponding layer that allows DTRs to capture characteristics of histology images properly, we have evaluated the accuracy of DTRs from various representative pre-trained models to capture similarity between histopathology images.

First, 21 images for each of the 32 cancer types were randomly selected from the preprocessed 271,700 patches (128 × 128 µm), of which one image per cancer type was chosen as a query. From the remaining 640 images, three trained pathologists independently selected three images that were histologically similar to each query, and ranked them according to the similarity to the query image.

We used DTRs for each of the representative convolutional layers of 6 common CNN architectures VGG19 (Simonyan and Zisserman, 2014), MobileNet V2 (Sandler et al., 2018), Inception V3 (Szegedy et al., 2015), NasNet A Large (Zoph et al., 2017), DenseNet (Huang et al., 2016) and Inception ResNet V2 (Szegedy et al., 2015)). First, cosine similarities between the q^th^ query image or the one rotated by 90 degrees and the selected i^th^ image were calculated, and the larger value was retained as *s*_*q,i*_.

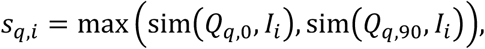

where *Q*_*q*,0_ is a q^th^ query image, *Q*_*q*,90_ is a q^th^ query image rotated 90 degrees, *I* is an evaluated image, and sim is a cosine similarity function.

Here, we denote a vector composed of *s*_*q,i*_ by *S*_*q*_.

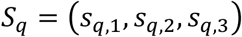

Then the vector of Spearman rank correlation coefficient with the rank by a trained pathologist was calculated as follows:

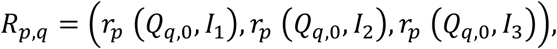

where *r*_*p*_ is a rank assigned by the p^th^ pathologist. Finally, the mean correlation over the three pathologists and query images were used as the evaluation index;

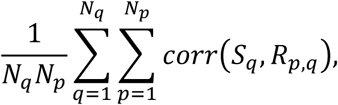

where *N*_*q*_ is the total number of query images (=32) and *N*_*p*_ is the total number of trained pathologists (=3). DTRs were implemented using Keras version 2.2.4 with Tensorflow 1.10.0 backend.

We also compared the performance of DTRs with other traditional texture features, histogram of oriented gradients (HOGs) and local binary pattern (LBP). We used HOG implemented in https://github.com/pochih/CBIR with various parameters (type = “global” and type-local” with n_slice = 2, 4, 8 and 16), and histogram of LBP implemented in scikits-image (Walt et al., 2014) with various radii (from 2–7). Kullback-Leibler divergence was used to measure similarity.

### Visualization of DTR distributions

the nonlinear dimension reduction algorithm Uniform Manifold Approximation and Projection (UMAP) (Mclnnes et al., 2018) was used to visualize DTRs. The averaged DTRs in each case or region were used so that one patient or region would occupy one point in the 2D space. Python’s UMAP library (https://github.com/lmcinnes/umap) was used for calculation with n_neighbors=10, min_dist=0.2 and metric-’cosine distance” for the visualization of a pan-cancer dataset. We used different parameters for the visualization of specific cancer types, as the data size varied for data of different cancer types, for example: n_neighbors = 13, and min_dist = 0.8 for prostate adenocarcinoma (PRAD) and stomach adenocarcinoma (STAD); n_neighbors = 7 and min_dist = 0.4 for pancreas adenocarcinoma (PAAD); and n_neighbors = 13 and min_dist = 0.4 for thyroid carcinoma (THCA). The Gleason score for PRAD and Lauren classification for STAD in each region were evaluated by trained pathologists.

### Color normalization

Color normalization was performed using the method of Reinhard et al. (2001) implemented in StainTools (https://github.com/Peter554/StainTools). Reference patches were selected from cancerous regions in TCGA dataset by a pathologist (Table S8), and color normalizations were performed separately for each cancer type.

### Genomic aberration and clinical data of TCGA

#### Somatic mutations

For cancer driver genes, actionable alterations and all alterations (driver alterations) were downloaded from the OncoKB database (Chakravarty et al., 2017) of the Memorial Sloan Kettering Cancer Center. There were 44 and 322 genes found for actionable and driver alterations, respectively. SNVs, indels, CNVs and gene fusions were used for analyses. TCGA cases with each driver mutation were retrieved from ‘tcga pan can atlas 2018’ database using the “cgdsr” Bioconductor package. The modified script of ‘AnnotatorCore.py’ of oncokb-annotator (https://github.com/oncokb/oncokb-annotator) was used to classify the mutations into actionable or driver alterations, as well as others. TCGA cases with fusion genes were downloaded from Tumor Fusion Gene Data Portal (http://www.tumorfusions.org/) developed by the US Jackson Laboratory. Driver genes mutated in at least three patients for at least one of the cancer types were used for evaluation, which resulted in 208 annotated and 30 actionable genes.

#### Mutational signatures

Forty-seven SBS mutational signatures identified by SigProfiler in a previous study (Alexandrov et al., 2020) were analyzed. Several SBS signatures with similar etiologies were merged (Table S9). For each SBS signature, positive cases were defined as having at least one mutation derived from the signature. Only those signatures found to be positive in more than two patients with at least one cancer type were used for the evaluation, which resulted in 28 SBS signatures.

#### BRCAness genomic signature

Based on a previous paper (Riaz et al., 2017), we defined BRCAness as being positive when a case had the largest number of signature 3 mutations among all signatures, and the number of large-scale state transitions (LST) was ≥ 15. Contribution of each mutational signature and LST information was downloaded from https://github.com/riazn/biallelic_hr.

#### Microsatellite instability

Definition of cases with MSI or defined as being microsatellite stabile (MSS) for STAD and COAD/READ were taken from prior research (Kather et al., 2019). Briefly, patients who were previously defined as MSI-High (MSI-H) or patients with a mutation count of >1,000 were defined as having MSI. The MSI/MSS label for each patient was downloaded from the corresponding Zenodo repository (Kather, 2019).

#### Histological subtype

Histological subtypes of the TCGA cases were downloaded from National Cancer Institute (NCI) Genomic Data Commons (https://gdc.cancer.gov/about-data/publications/pancanatlas) on November 12^th^, 2018. The term ‘histological_type’ in clinical_PANCAN_patient_with_followup.tsv was used for query.

### Prediction of genomic features from histopathology images using DTRs and supervised machine learning

The presence of a driver gene mutation or mutation signature of each cancer type was predicted using logistic regression, where 1,024-dimensional DTRs from histopathology images were used as features. Stratified 3-fold cross validation for the cases—and not for images, as to avoid patches from the same case being contained in both training and test data— was performed for each type of mutation and cancer. During the training phase, all the patches were treated independently irrespective of the derived cases, but the prediction as well as AUC score calculation were performed for each case by averaging the probability score of the logistic regression model for all the patches from the case. The prediction was made for six magnification levels (128-256 µm), and the optimal model was selected. AUC scores from 100,000 random permutations were also calculated and compared to the median AUC in 3-fold cross-validation to calculate the p-value. Finally, for the genomic aberrations with mean AUC > 0.65, the logistic regression models were trained using all the available data at three magnification levels (128, 192, and 256 µm) and implemented into the web system.

The programming language Python version 3.6.3 and the machine learning library scikit-learn version 0.20.0 were used for logistic regression, cross validation, and AUC calculation (color norm, clinical). For logistic regression, SGD Classifier with loss = ‘log’, penalty = ‘l2’, alpha = 0.0001 was applied.

### External validation cohort of genomic aberration prediction

#### Informed consent and sample preparation

Fresh frozen gastric cancer and paired normal stomach tissues were obtained from patients who underwent gastrectomy at the University of Tokyo Hospital (n = 91). Informed consent was obtained from each subject, and this study was approved by the institutional review boards at the University of Tokyo. Genomic DNA was extracted using QIAamp DNA Mini kit (Qiagen, Hilden, Germany) according to the manufacturer’s instructions.

#### Whole-exome sequencing and mutation analysis

Whole-exome sequencing was performed for 91 gastric cancers and paired normal gastric tissues as previously described (Kakiuchi et al., 2014). Each DNA sample (1.1 µg) was sheared using a Covaris SS Ultrasonicator (Covaris, MA, USA) as per the manufacturer’s instructions. A Sciclone NGS workstation (Caliper Life Sciences, MA, USA) was used to automatically construct the DNA sequence libraries according to the manufacturer’s instruction. Exome regions were captured by an Agilent SureSelect Human All Exon V4 Kit and V5+ LincRNA (Agilent Technologies, CA, USA). Sequencing was performed on an Illumina HiSeq 2000 instrument (Illumina, CA, USA) and the provided protocol for producing 2 × 100-bp paired-end reads. Image analyses and base calling were performed using the Illumina pipeline with default settings.

The reads were aligned to the human reference genome GRCh37/hg19 using the Burrows-Wheeler Aligner (BWA) and NovoAlign software (Novocraft Technologies, Selangor, Malaysia). After removal of PCR duplicates, the Short-Read Micro re-Aligner (SRMA) method was used to improve variant discovery via local realignments (Homer and Nelson, 2010). Variant calling was performed using an integrated genotyping software (karkinos: http://github.com/genome-rcast/karkinos) with default settings as previously reported (Kakiuchi et al., 2014).

#### Prediction of somatic mutations from histopathology images using DTRs

H&E stained WSIs from 91 gastric cancer tissues with somatic mutation information were used to validate the classification model for gene mutation prediction from histopathology images using DTRs. Slides were digitized using a Hamamatsu Nanozoomer 2.0 HT whole slide scanner. For each of the resulting whole slide images in Hamamatsu ndipi format, at least three representative cancerous regions were selected as polygons by a trained pathologist as performed within the TCGA dataset using the same criteria. Then, 10 patches sized 128 × 128 µm were arbitrarily cropped using random angles from each of the annotated region using keras-OpenSlideGenerator, and were then resized to 256 × 256 pixels and input to the mutation predictors described above. *TP53, CDH1*, and *PTEN* genes were mutated in > five patients and thus were chosen for validation. AUC score was calculated for each case by averaging the probability score of the logistic regression model for all the patches from the same case.

### Visualization using Grad-CAM

The Grad-CAM algorithm implemented in pytorch (https://github.com/1Konny/gradcam_plus_plus-pytorch) was used to visualize the relevant regions in histology images for the prediction of genomic aberrations. Because Grad-CAM requires fine-tuned CNN models for accurate visualization, we fine-tuned the VGG16 network pre-trained on ImageNet for predicting *TP53* mutations in BRCA, MSI status in STAD, *IDH1* mutations in GBM and *U2AF1* mutations in LUAD, all of which are predictable by the logistic regression model using DTRs. All cases were randomly split into training, validation, and test datasets with a 4:1:1 ratio. The network was trained using Stochastic Gradient Descent optimizer with learning rate = 0.00005 and momentum = 0.99. In the first three epochs, only the fully-connected layers were trained, and in the remaining 12 epochs, all layers were trained. The model with minimum validation loss was used for GradCAM. Pytorch 1.0 was used to train the network.

### Luigi system for CBIR and genomic aberration prediction

#### Implementation

The Luigi system was developed with a Python backend and Clojure frontend; thus, it can be accessed by any modern web browser on Mac OS X or Windows operating system. Users can search images relevant to a query image from the database, which is comprised of 475,701 patches representing 32 cancer types from TCGA according to the aforementioned selection procedure using intuitive operations. Links for the cBioPortal (Cerami et al., 2012) website are also provided, enabling the users to access clinical, pathological, and molecular information relevant to the retrieved case. The Luigi system can be used without registration, but if a user account is created, the system will save the entire image upload history. Although the Luigi database contains WSI information for each case, the query and the retrieved images handled by the system are patches extracted from WSIs because, in most situations, users submit small regions of a whole slide with specific features.

#### CBIR and prediction of genomic aberrations

When a query image is submitted to the Luigi system, 1,024-dimensional DTRs of the image, as well as the one image rotated 90° for imposing rotational invariance, are calculated. The system then searches for the most similar images in the database using the cosine similarity of the extracted DTRs between the query and database images. An approximate nearest neighbor search method with the randomized kd-tree algorithm implemented in pyflann (Muja and Lowe, 2009) was adopted to enable high-speed searching. For the prediction of genomic aberrations, the magnification level of the query image was first estimated by a Support Vector Machine classifier trained on the DTRs of cancerous regions for each cancer type in TCGA images at three magnification levels (128, 192, and 256 µm). Subsequently the logistic regression classifier trained at the predicted magnification level was applied for each genomic aberration.

#### User interface of the Luigi web system

Figure 7A shows the interface of the Luigi system. Users upload a histopathological image file by either dragging and dropping the file, pasting a screenshot of the image, or selecting “click here to select a file.” The uploaded image is resized to 256 × 256 pixels (the size of the images in the database). Because this resizing process could alter the texture, uploading images with appropriate magnification levels is essential to retrieve good results. Then, the Luigi system will retrieve the most relevant patch in each WSI from all cancer types, rank them according to cosine similarity and display them. Tissue origin can be changed using the pull-down menu above the results. When a user selects any of the retrieved images, that image will become a new query and can be used to retrieve relevant patches by clicking the “Search” button. The TCGA slide ID (TCGA-XX-XXXX-XXX-XX-XXX) accompanying each image contains a link to the cBioPortal website (Cerami et al., 2012), so users can open a new window showing the summary of the selected case. There users can review the pathology report as well as clinical and molecular information for the case. Predicted genomic aberrations are shown in the right panel. The result from the estimated cancer type, i.e. the most frequent type found within the top 10 retrieved cases, is initially shown, but users can alter the cancer type manually. Luigi is available at https://luigi-pathology.com/

#### Luigi application

A smartphone application was also developed to allow users to submit appropriately scaled query images directly to the Luigi web application and view retrieved images. Figure 7C displays this interface. Users can take a photo of their cases or select an image stored in the smartphone, and then adjust the query image size while referring to the image on top. Users can then press the check button to display the relevant cases to the uploaded query image found for all tumors with the order of cosine similarity. The tissue of origin can be changed by selecting from the drop-down menu at the top of the screen. Users can also access to the cBioPortal website by tapping TCGA slide ID of the retrieved case. An iOS version of the smartphone application is currently available.

#### CBIR performance evaluation

We evaluated the performance of Luigi web systems to retrieve similar images in the following 12 cancer types: lower grade glioma (LGG), thyroid carcinoma (THCA), kidney renal clear cell carcinoma (KIRC), lung adenocarcinoma (LUAD), breast cancer (BRCA), colon adenocarcinoma (COAD), head and neck squamous cell carcinoma (HNSC), kidney chromophobe (KICH), thymoma (THYM), prostate adenocarcinoma (PRAD), sarcoma (SARC), and testicular germ cell tumor (TCGT). The query images representing typical morphology were selected from TCGA images by a trained pathologist, and retrieved images from all the cancer types were evaluated. We evaluated precision based on the cancer type except for LUAD, COAD, HNSC, in which adenocarcinoma or squamous cell carcinoma were used as the ground truth category. Precision values for the top 5, 10, and 20 images were evaluated. In another analysis, we prepared query images for 12 cancer types. We then created a dataset for each query containing 19 randomly selected images and the most relevant image selected by the Luigi. Five trained pathologists were asked to blindly select those most histologically similar to the query image, irrespective of cancer types.

Query images for the Luigi smartphone application were taken with a microscope mounted on an iPhone 8 using a Gosky Universal Cell Phone Adapter Mount (American Gosky Optics, GA, USA. http://www.goskyoptics.com/). We have selected 5 tumor regions from within our own WSIs—including human stomach adenocarcinoma and patient-derived tumor xenograft tissues from colorectal and pancreas adenocarcinoma—and used these as queries. The magnification levels were adjusted manually.

### Estimation of the number of patients with predictable actionable mutations

We have roughly estimated the number of patients worldwide whose actionable mutations can be predicted solely via histology images. The number of incidences of each cancer type were downloaded from the Cancer Today website from the World Health Organization (https://gco.iarc.fr/today/online-analysis-map). The downloaded data contained crude incidence rates and were labeled with cancer types. Because some of the labeled cancer sites did not correspond to those of the TCGA’s, additional processing was necessary. We have combined “lip, oral cavity,” “hypopharynx,” “larynx,” “nasopharynx,” and “oropharynx” cancer sites as being under the category of “HNSC.” Because GBM and LGG in TCGA fall into “brain, central nervous system” for cancer sites, we estimated the incidence of each of these cancers using the average incidence rate of nervous system and intracranial tumors in England (Maile et al., 2016). For lung cancer, international incidence rates of lung adenocarcinoma and squamous cell carcinoma within all lung cancer types (Cheng et al., 2016) were used for estimation. Some of the cancer types in TCGA—such as TCGT, THYM, UCS, and SARC—that were not included in the WHO analysis, were not used due to a lack of statistics.

The latest available population data (mostly from 2018) in each country was downloaded from The World Bank (**https://data.worldbank.org/**).

We used actionable mutations with a positive predictive value (PPV) > 0.7 for the estimation. PPVs for each gene in specific cancer types were estimated from TCGA population as follows: (1) when PPVs based on an optimal classification threshold, which was determined by the Youden index (Youden, 1950) for the mean ROC curve among 3-fold cross-validations, exceed 0.7, the PPV and the corresponding true positive rate (TPR) was used. (2) Otherwise, when the maximum PPV in all the classification thresholds exceeded 0.7, the PPV and the corresponding TPR was used. The number of patients with predictable actionable mutations was estimated by the number of cancer incidence multiplied by the true positive rate of the mutation in TCGA population calculated for each of the corresponding PPVs. Table S7 shows the selected genes. The total number of patients who can be found to be actionable mutation-positive using our method was calculated by summing up all the estimated number of patients for each actionable gene or genomic aberration.

## Tables

**Table S1. Number of patients and diagnostic WSIs for each cancer type used in this study, Related to Figure 1.**

**Table S2. Predictable annotated driver mutations (q < 0.02), Related to Figure 4.**

**Table S3. Predictable actionable driver mutations (q < 0.02), Related to Figure 4.**

**Table S4. Predictable signatures (q < 0.02), Related to Figure 4.**

**Table S5. Other predictable genomic features (q < 0.02), Related to Figure 4.**

**Table S6. classic histopathologic keywords for the morphologic features in mutant-positive cases, Related to Figures 5 and 6.**

**Table S7. Estimated Positive Prediction Values and True Positive Rate for actionable mutation prediction, Related to Figure 7.**

**Table S8. Reference WSIs used for color normalization, Related to Methods.**

**Table S9. Mutational signatures analyzed in this study, Related to Methods.**

## Videos

**Supplementary Video 1. Usage of smartphone application, Related to Figure 7.**

## Lead Contact

Further information and requests for resources should be directed to and will be fulfilled by the Lead Contact, Shumpei Ishikawa.

## Acknowledgements

The slide images and the corresponding cancer information were downloaded from the Genomic Data Commons portal (https://gdc-portal.nci.nih.gov) and are in whole or in part based upon data generated by the TCGA Research Network (http://cancergenome.nih.gov/). We thank Enago (www.enago.jp) for the English language review. We thank Ruri Saito, Munetoshi Hinata, Hirofumi Rokutan, Naohiro Makise, and Sho Yamazawa for evaluating a system. We also thank Menghua Zhang for helping us take a Supplementary Video. This study was supported by the AMED Practical Research for Innovative Cancer Control under Grant Number JP 19ck0106400 to S.I.

## Author Contributions

Conceptualization, S.I. and D.K.; Methodology, K.F. and D.K. ; Software, D.K., K.T., R.S., Hiroki Konishi and Toshikazu Umezaki.; Validation, D.K.; Formal Analysis, D.K. and K.S. ; Investigation, D.K. and S.I., Hirotomo Koda, G.F., S.N., Hiroto Katoh, and A.K.; Resources, D.K, Tetsuo Ushiku and M.F.; Data Curation, Hirotomo Koda and A.K.; Writing - Original Draft D.K. and S.I.; Writing - Review & Editing, D.K. and S.I. ; Visualization, D.K. F.M.; Supervision, S.I.

## Declaration of interests

The authors declare no competing interests

## Notes

### Competing Interest Statement

The authors have declared no competing interest.

## References

Alexandrov, L.B., Kim, J., Haradhvala, N.J., Huang, M.N., Tian Ng, A.W., Wu, Y., Boot, A., Covington, K.R., Gordenin, D.A., Bergstrom, E.N., et al. (2020). The repertoire of mutational signatures in human cancer. Nature 575, 94–101.

Baena, E., Gandarillas, A., Vallespinos, M., Zanet, J., Bachs, O., Redondo, C., Fabregat, I., Martinez-A, C., and de Alboran, I.M. (2005). c-Myc regulates cell size and ploidy but is not essential for postnatal proliferation in liver. Proc. Natl. Acad. Sci. U. S. A. 102, 7286–7291.

Bass, A.J., Thorsson, V., Shmulevich, I., Reynolds, S.M., Miller, M., Bernard, B., Hinoue, T., Laird, P.W., Curtis, C., Shen, H., et al. (2014). Comprehensive molecular characterization of gastric adenocarcinoma. Nature 513, 202–209.

Bejnordi, B.E., Veta, M., Diest, P.J. van, Ginneken, B. van, Karssemeijer, N., Litjens, G., Laak, J.A.W.M. van der, Hermsen, M., Manson, Q.F., Balkenhol, M., et al. (2017). Diagnostic assessment of deep learning algorithms for detection of lymph node metastases in women with breast cancer. JAMA 315, 21992210.

Cancer Genome Atlas Research Network, Brat, D.J., Verhaak, R.G.W., Aldape, K.D., Yung, W.K.A., Salama, S.R., Cooper, L.A.D., Rheinbay, E., Miller, C.R., Vitucci, M., et al. (2015). Comprehensive, integrative genomic analysis of diffuse lower-grade gliomas. N. Engl. J. Med. 372, 2481–2498.

Cerami, E., Gao, J., Dogrusoz, U., Gross, B.E., Sumer, S.O., Aksoy, B.A., Jacobsen, A., Byrne, C.J., Heuer, M.L., Larsson, E., et al. (2012). The cBio Cancer Genomics Portal: an open platform for exploring multidimensional cancer genomics data. Cancer Discov. 2, 401–404.

Chakravarty, D., Gao, J., Phillips, S., Kundra, R., Zhang, H., Wang, J., Rudolph, J.E., Yaeger, R., Soumerai, T., Nissan, M.H., et al. (2017). OncoKB: A Precision Oncology Knowledge Base. JCO Precis. Oncol. 1–16.

Cheng, T.-Y.D., Cramb, S.M., Baade, P.D., Youlden, D.R., Nwogu, C., and Reid, M.E. (2016). The international epidemiology of lung cancer: latest trends, disparities, and tumor characteristics. J. Thorac. Oncol. Off. Publ. Int. Assoc. Study Lung Cancer 11, 1653–1671.

Coudray, N., Ocampo, P.S., Sakellaropoulos, T., Narula, N., Snuderl, M., Fenyo, D., Moreira, A.L., Razavian, N., and Tsirigos, A. (2018). Classification and mutation prediction from non-small cell lung cancer histopathology images using deep learning. Nat. Med. 24, 1559.

Dalal, N., and Triggs, B. (2005). Histograms of oriented gradients for human detection. In 2005 IEEE Computer Society Conference on Computer Vision and Pattern Recognition (CVPR’05), pp. 886–893 vol. 1.

Deng, J., Dong, W., Socher, R., Li, L., Li, K., and Fei-fei, L. (2009). Imagenet: A large-scale hierarchical image database. In 2009 IEEE Conference on Computer Vision and Pattern Recognition, p.

Gao, Y., Beijbom, O., Zhang, N., and Darrell, T. (2015). Compact Bilinear Pooling. 151106062 Cs.

Hanahan, D., and Weinberg, R.A. (2011). Hallmarks of cancer: the next generation. Cell 144, 646–674.

Hatipoglu, N., and Bilgin, G. (2016). Feature extraction for histopathological images using Convolutional Neural Network. In 2016 24th Signal Processing and Communication Application Conference (SIU), pp.645–648.

Homer, N., and Nelson, S.F. (2010). Improved variant discovery through local re-alignment of short-read next-generation sequencing data using SRMA. Genome Biol. 11, R99.

Huang, G., Liu, Z., van der Maaten, L., and Weinberger, K.Q. (2016). Densely Connected Convolutional Networks. 160806993 Cs.

Kakiuchi, M., Nishizawa, T., Ueda, H., Gotoh, K., Tanaka, A., Hayashi, A., Yamamoto, S., Tatsuno, K., Katoh, H., Watanabe, Y., et al. (2014). Recurrent gain-of-function mutations of RHOA in diffuse-type gastric carcinoma. Nat. Genet. 46, 583–587.

Kather, J.N. (2019). Histological images for MSI vs. MSS classification in gastrointestinal cancer, FFPE samples (Zenodo).

Kather, J.N., Pearson, A.T., Halama, N., Jager, D., Krause, J., Loosen, S.H., Marx, A., Boor, P., Tacke, F., Neumann, U.P., et al. (2019). Deep learning can predict microsatellite instability directly from histology in gastrointestinal cancer. Nat. Med. 25, 1054–1056.

Lin, T.-Y., RoyChowdhury, A., and Maji, S. (2015). Bilinear CNN Models for fine-grained visual recognition. 150407889 Cs.

Maile, E.J., Barnes, I., Finlayson, A.E., Sayeed, S., and Ali, R. (2016). Nervous System and Intracranial Tumour Incidence by Ethnicity in England, 2001-2007: A Descriptive Epidemiological Study. PLOS ONE 11, e0154347.

Masaki, T. (2006). Tumor budding in colorectal cancer: recent progress in colorectal cancer research (Nova Publishers).

McInnes, L., Healy, J., and Melville, J. (2018). UMAP: Uniform Manifold Approximation and Projection for Dimension Reduction. 180203426 Cs Stat.

Muja, M., and Lowe, D.G. (2009). Fast approximate nearest neighbors with automatic algorithm configuration. In VISAPP International Conference on Computer Vision Theory and Applications, pp. 331–340.

Nagpal, K., Foote, D., Liu, Y., Po-Hsuan, Chen, Wulczyn, E., Tan, F., Olson, N., Smith, J.L., Mohtashamian, A., et al. (2018). Development and validation of a deep learning algorithm for improving Gleason scoring of prostate cancer.

Ojala, T., Pietikainen, M., and Maenpaa, T. (2002). Multiresolution gray-scale and rotation invariant texture classification with local binary patterns. IEEE Trans. Pattern Anal. Mach. Intell. 24, 971–987.

Pharoah, P.D.P., Guilford, P., and Caldas, C. (2001). Incidence of gastric cancer and breast cancer in CDH1 (E-cadherin) mutation carriers from hereditary diffuse gastric cancer families. Gastroenterology 121, 13481353.

Phillips, S.M., Banerjea, A., Feakins, R., Li, S.R., Bustin, S.A., and Dorudi, S. (2004). Tumour-infiltrating lymphocytes in colorectal cancer with microsatellite instability are activated and cytotoxic. Br. J. Surg. 91, 469–475.

Reinhard, E., Adhikhmin, M., Gooch, B., and Shirley, P. (2001). Color transfer between images. IEEE Comput. Graph. Appl. 21, 34–41.

Riaz, N., Blecua, P., Lim, R.S., Shen, R., Higginson, D.S., Weinhold, N., Norton, L., Weigelt, B., Powell, S.N., and Reis-Filho, J.S. (2017). Pan-cancer analysis of bi-allelic alterations in homologous recombination DNA repair genes. Nat. Commun. 5, 1–7.

Sandler, M., Howard, A., Zhu, M., Zhmoginov, A., and Chen, L.-C. (2018). MobileNetV2: Inverted Residuals and Linear Bottlenecks. 180104381 Cs.

Schaumberg, A.J., Rubin, M.A., and Fuchs, T.J. (2017). H&amp;E-stained whole slide image deep learning predicts SPOP mutation state in prostate cancer. BioRxiv 064279.

Simonyan, K., and Zisserman, A. (2014). Very deep convolutional networks for large-scale image recognition. ArXiv Prepr. 14091556.

Szegedy, C., Vanhoucke, V., Ioffe, S., Shlens, J., and Wojna, Z. (2015). Rethinking the Inception Architecture for Computer Vision. 151200567 Cs.

Walt, S. van der, Schonberger, J.L., Nunez-Iglesias, J., Boulogne, F., Warner, J.D., Yager, N., Gouillart, E., and Yu, T. (2014). scikit-image: image processing in Python. PeerJ 2, e453.

Weinstein, J.N., Collisson, E.A., Mills, G.B., Shaw, K.R.M., Ozenberger, B.A., Ellrott, K., Shmulevich, I., Sander, C., Stuart, J.M., Network, C.G.A.R., et al. (2013). The cancer genome atlas pan-cancer analysis project. Nat. Genet. 45, 1113–1120.

Yosinski, J., Clune, J., Nguyen, A., Fuchs, T., and Lipson, H. (2015). Understanding Neural Networks Through Deep Visualization. 150606579 Cs.

Youden, W.J. (1950). Index for rating diagnostic tests. Cancer 3, 32–35.

Zanet, J., Pibre, S., Jacquet, C., Ramirez, A., Alboran, I.M. de, and Gandarillas, A. (2005). Endogenous Myc controls mammalian epidermal cell size, hyperproliferation, endoreplication and stem cell amplification. J. Cell Sci. 115, 1693–1704.

Zoph, B., Vasudevan, V., Shlens, J., and Le, Q.V. (2017). Learning Transferable Architectures for Scalable Image Recognition.

