## Supplementary Figures and Tables for "Deep Texture Representations as a Universal Encoder for Pan-cancer Histology"

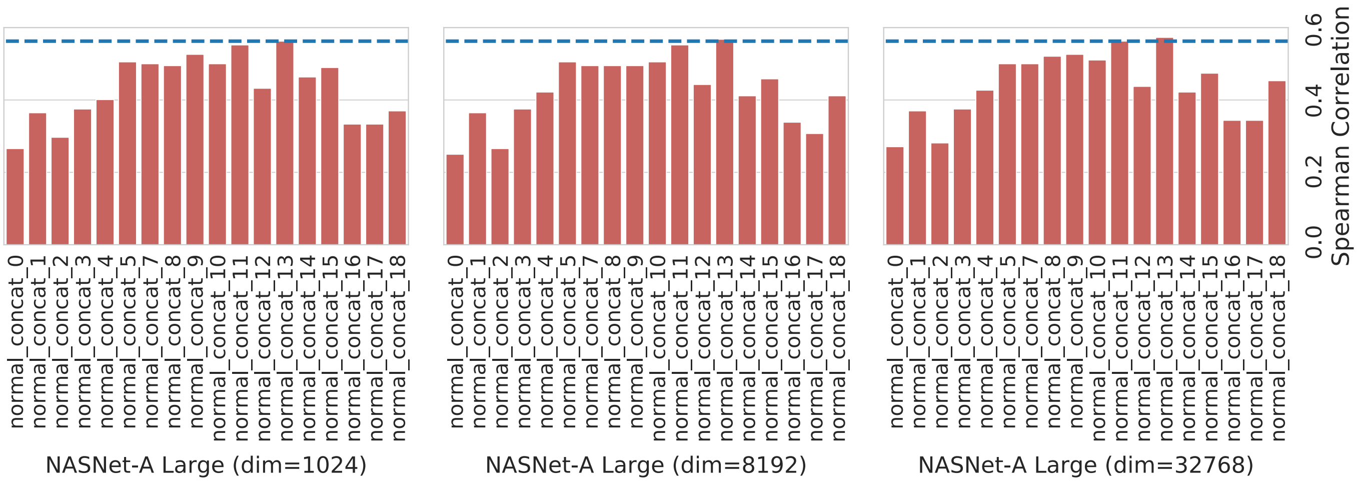


**Figure S1 Evaluation of DTRs for multiple dimensions, Related to Figure 1C.**

Spearman's rank correlation between the similarity of the images ranked by pathologists and by 1024-, 8192-, or 32768-dimensional DTRs from the 'normal_concat_11' layer in the NASNet-A Large network. The layers are deeper along the x-axis from left to right in each network.

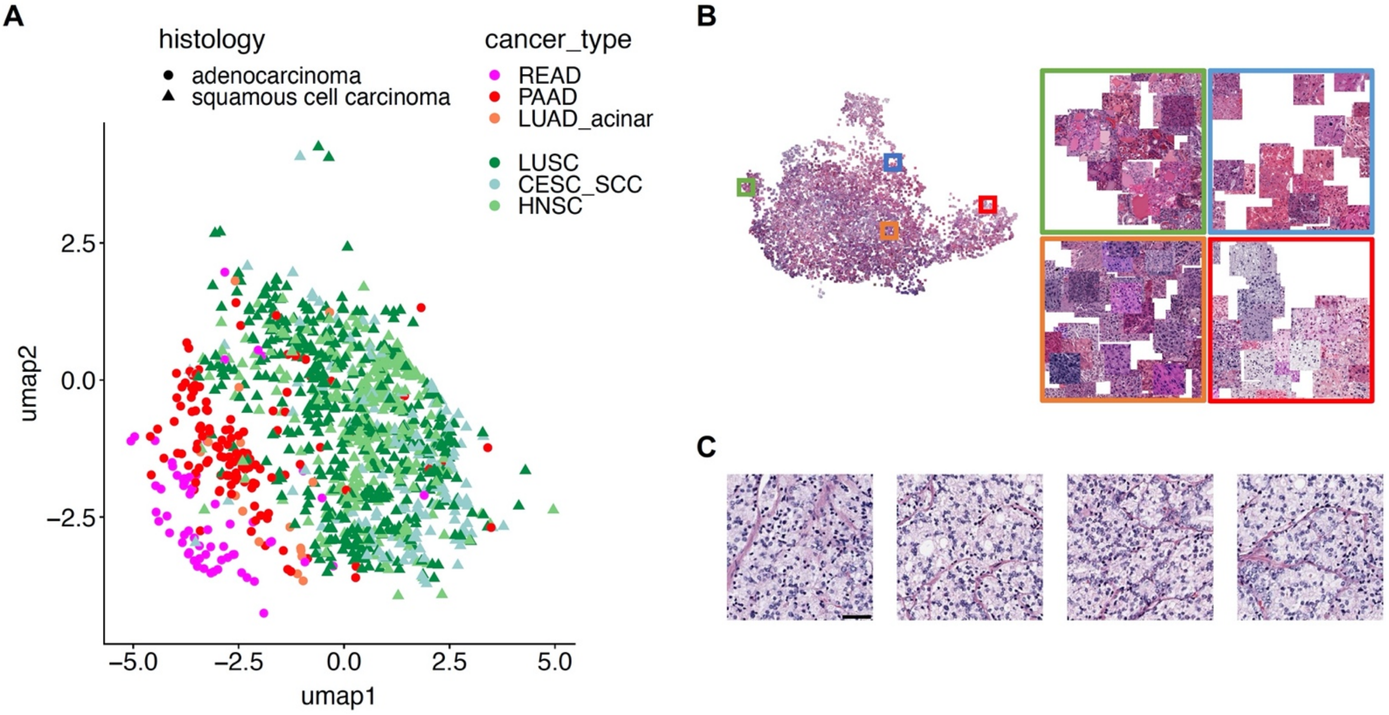


**Figure S2 Encoding histological features by DTR, Related to Figure 2.**

(A) Distribution of adenocarcinomas and squamous cell carcinomas of various organs in UMAP projection.

(B) 2D projection of the DTRs of TCGA cancer histology images from Figure 2. Enlarged view of the boxed regions with corresponding frame color shows that histologically similar images—regardless of the staining intensity/color variation—are grouped close together.

(C) A case exhibiting hypernephromatoid pattern of prostate adenocarcinoma (TCGA patient ID: TCGA-WW-A8ZI), which were indicated with arrows in Figure 2C. Scale bar: 50 μm.

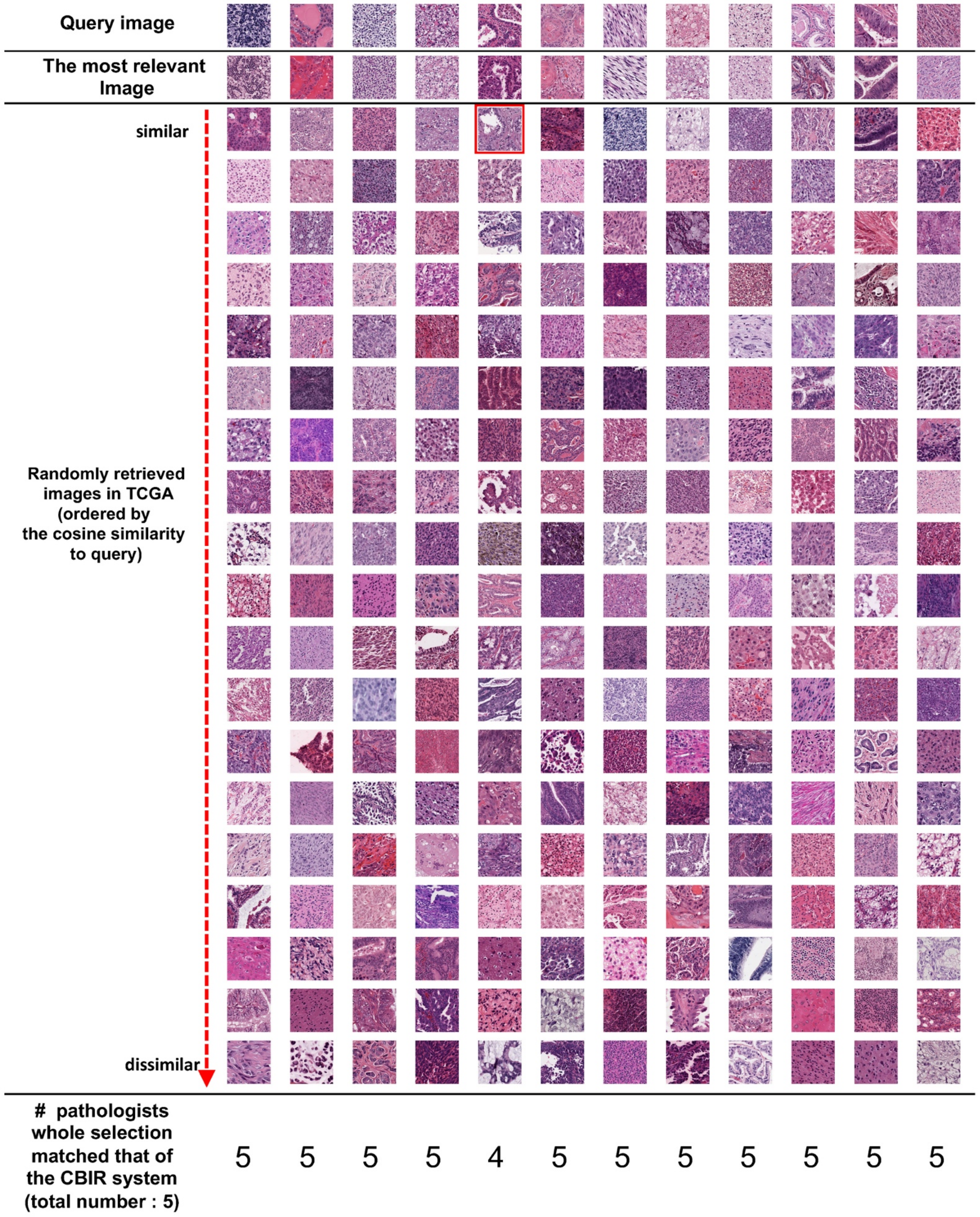


**Figure S3 Blinded comparison between similar images retrieved by DTR-based CBIR and histopathologically similar images selected by trained pathologists, Related to Figure 3.**

From top to bottom, 12 series of representative query images from various cancer types, with the most relevant image to each query selected by the CBIR system, and 19 randomly selected images from TCGA are shown. Out of a total of 20 images—including the one selected by the CBIR system—five trained pathologists selected the image they judged to be the most similar morphologically in a strictly blinded manner. The bottom row indicates the number of pathologists whose selection matched that of the CBIR system. The image framed in red was selected by a pathologist, but did not match the image selected by the CBIR system.

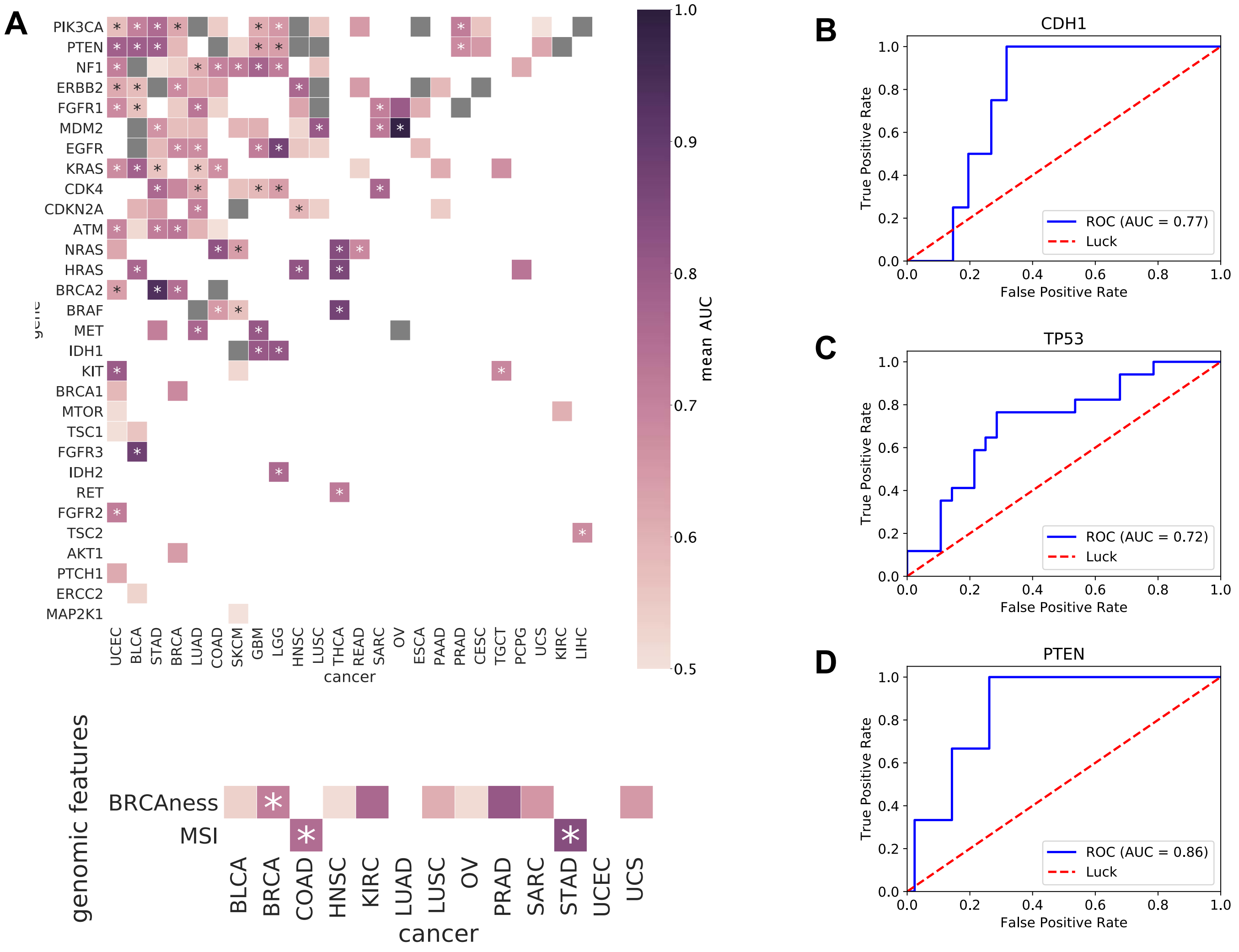


**Figure S4 Clinically actionable somatic gene mutations and genomic features predicted from histology images, Related to Figure 4.**

(A) Area under the receiver operating characteristic curve values for the prediction of somatic mutations of informative OncoKB clinically actionable gene mutations (30 out of 44), shown as in Figure 4A (top). BRCAness and MSI status are also shown (bottom). Asterisks indicate $q < 0.02$.

(B–D) Performance of the logistic regression models using DTRs trained with TCGA STAD datasets on our independent STAD cohort were measured in terms of AUC for *CDH1* (B), *TP53* (C), and *PTEN* (D) mutations.

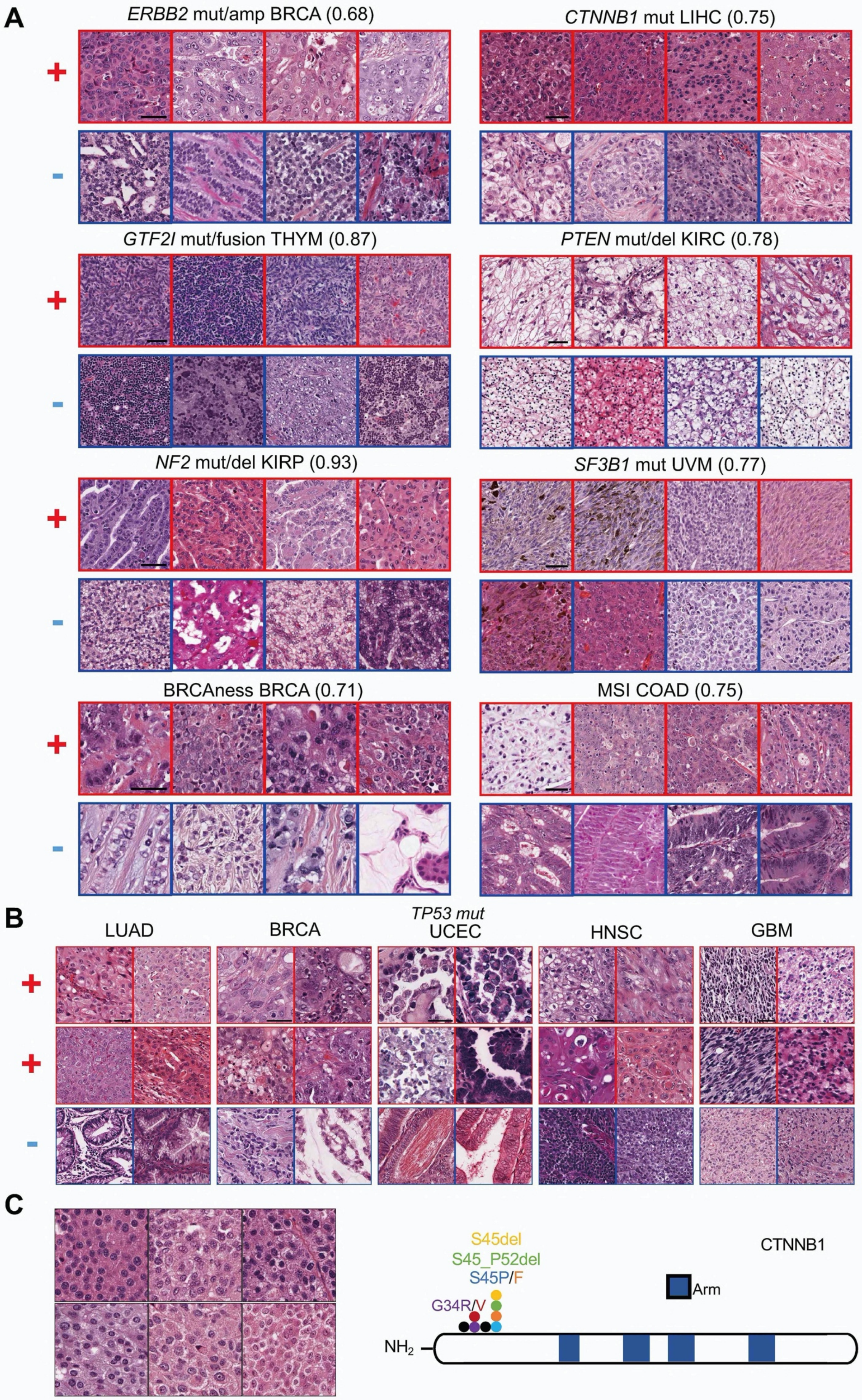
A
ERBB2 mut/amp BRCA (0.68)
CTNNB1 mut LIHC (0.75)
GTF2I mut/fusion THYM (0.87)
PTEN mut/del KIRC (0.78)
NF2 mut/del KIRP (0.93)
SF3B1 mut UVM (0.77)
BRCAness BRCA (0.71)
MSI COAD (0.75)
+
-
B
TP53 mut
LUAD
BRCA
UCEC
HNSC
GBM
C
S45del
S45_P52del
S45P/F
G34R/V
CTNNB1
Arm
$NH_2$

**Figure S5. Morphologic features correlated with genomic aberrations, Related to Figure 5.**

All images for each gene-cancer combination are shown in the same magnification. Scale bar: 50 µm.

(A) Representative images of cases with high (top row) and low (bottom row) somatic mutation probabilities are shown as in Figure 5: *ERBB2* in BRCA, *CTNNB1* in LIHC, *GTF2I* in THYM, *PTEN* in KIRC, *NF2* in KIRP, *SF3B1* in UVM, BRCAness in BRCA, and MSI in COAD (see Supplementary Methods).

(B) Common morphologic change, i.e. increased nuclear atypia and size, is observed in TP53-mutated LUAD, BRCA, UCEC, HNSC, and GBM. Scale bar: 50 µm.

(C) *CTNNB1*-mutated ACC cases exhibit similar histology to regularly packed tumor cells with round nuclei. *CTNNB1* mutations for all cases are concentrated in a hot spot region.

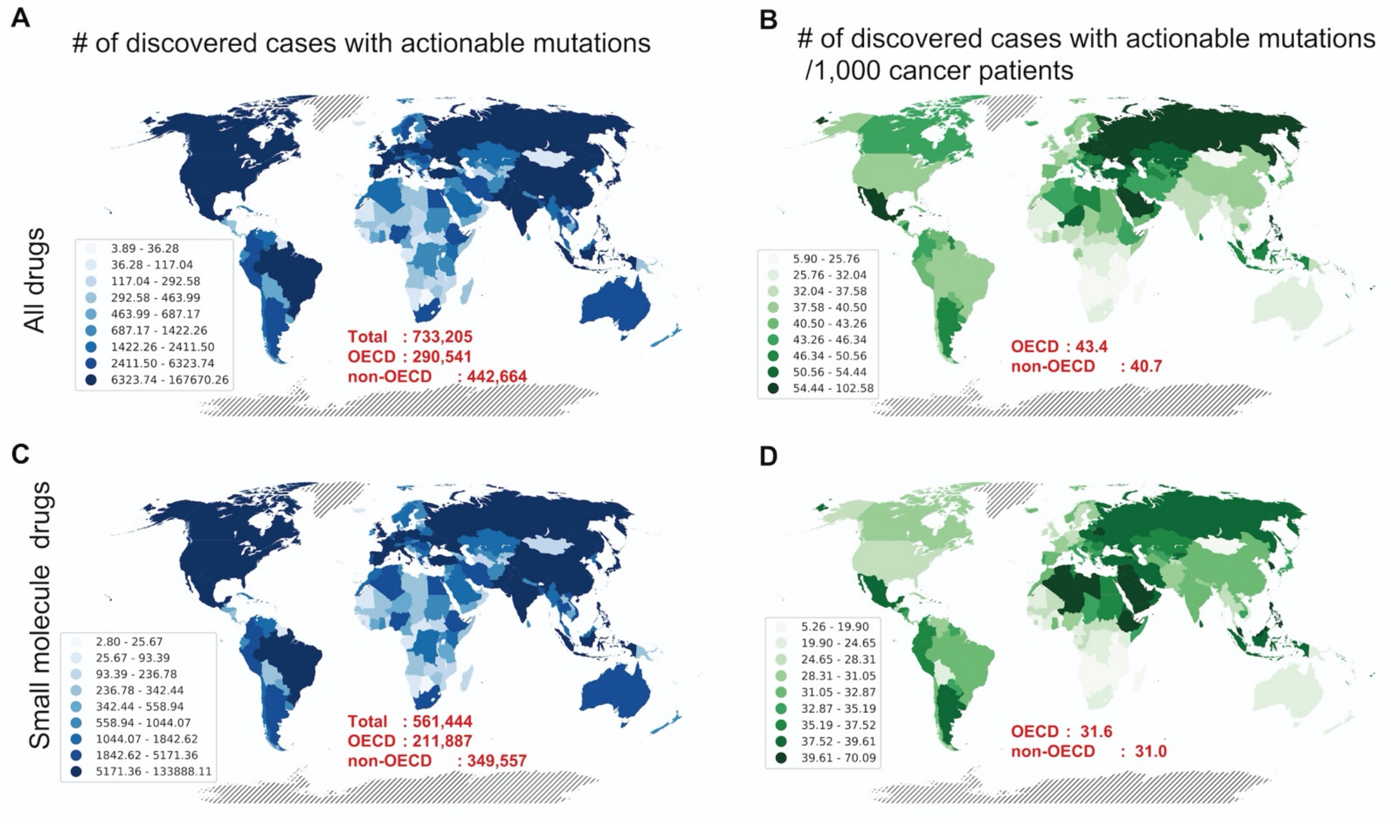


**Figure S6 Global impact of actionable mutation prediction from routine H&E histology images, Related to Figure 7D.**

Each country is shaded based on the estimated number of annual cancer cases with clinically actionable mutations that can be identified using our method with > 70% positive prediction value. (A) and (C) indicate the estimated numbers of identified cases at the theoretical maximum with mutations clinically actionable either by any drug or by small-molecule drugs, respectively. (B) and (D) show the rates of the identified cases per 1,000 cancer patients in each country that are clinically actionable by any drug or small-molecule drugs, respectively. Figure 7D is shown again here within (C) for comparison. The total number of cases (A and C) or the permil (B and D) numbers of people that can be found to be cancer-positive using our method in OECD and non-OECD countries annually are shown in red.

**Table S5. Other predictable genomic features (q < 0.02), Related to Figure 4.**

| cancer type | Genomic features | mean AUC | q-value |
|---|---|---|---|
| STAD | MSI-H | 0.837 | 0.00033 |
| COAD | MSI-H | 0.753 | 0.0041433 |
| BRCA | BRCAness | 0.710 | 0.00165 |

**Table S6. classic histopathologic keywords for the morphologic features in mutant-positive cases, Related to Figures 5 and 6.**

| Figure | Gene/Signature | Mutation type | Cancer-Type | descriptive histologic findings |
|---|---|---|---|---|
| 5A | FGFR3 | mut/amp/fusion | BLCA | regularly arranged cells with oval nuclei |
| 5A | IDH1 | mut | GBM | astrocyte-like faint or gemistocytic cytoplasm |
| 5A | TP53 | mut/del | BRCA | large atypical/pleomorphic nuclei |
| 5A | CTNNB1 | mut | SKCM | spindle-shaped cells with melanin pigment |
| 5A | BRCAness | signature | PRAD | poorly differentiated, without tube formation |
| 5A | MSI-H | signature | STAD | intra-epithelial lymphocytic infiltration |
| 5B | MYC | amp | LIHC/PAAD/LUSC/BRCA/ BLCA | increase in cell size with plump cytoplasm |
| 6D | U2AF1 | mut | LUAD | lepidic-predominant, with “hobnail” appearance |
| S5A | ERBB2 | mut/amp | BRCA | large nuclei with open chromatin |
| S5A | SF3B1 | mut | UVM | spindle-shaped cells with fascicular pattern |
| S5A | GTF2 | mut/fusion | THYM | small spindle-shaped cells with wavy nuclei |
| S5A | PTEN | mut/del | KIRC | mixture of irregular-shaped tumor cells and fibrous stroma |
| S5A | NF2 | mut/del | KIRP | abundant cytoplasm with conspicuous nucleoi |
| S5A | CTNNB1 | mut | LIHC | regular small nuclei without stromal reaction |
| S5A | BRCAness | signature | BRCA | discohesive clustering of cells with irregularly round nuclei |
| S5A | MSI-H | signature | COAD | intra-epithelial lymphocytic infiltration or mucinous type |
| S5B | TP53 | mut/del | LUAD/BRCA/UCEC/HNSC/GBM | large atypical/pleomorphic nuclei |
| S5C | CTNNB1 | mut | ACC | regularly packed cells with round nuclei |
| 6C | BRAF | mut/fusion | THCA | papillary structure with tall cells |
| 6C | RTKs | mut/fusion | THCA | cancerous(epithelial) front and back are reversed |
| 6C | RASs | mut | THCA | thyroid follicular structure |

**Table S7. Estimated Positive Prediction Values and True Positive Rate for actionable mutation prediction, Related to Figure 7.**

| Cancer type | gene | PPV | TPR | commonly used drugs |
|---|---|---|---|---|
| BLCA | FGFR3 | 0.77 | 0.0283 | Erdafitinib |
| BLCA | BRCAness | 0.70 | 0.0222 | platinum drug |
| BRCA | PIK3CA | 0.76 | 0.0209 | Buparlisib, Serabelisib, Copanlisib |
| COAD | KRAS | 0.72 | 0.2736 | Cobimetinib, Binimetinib, Trametinib |
| GBM | EGFR | 0.72 | 0.3645 | Erlotinib, Afatinib, Osimertinib |
| LGG | IDH1 | 0.90 | 0.5607 | Ivosidenib |
| OV | BRCAness | 0.86 | 0.4857 | platinum drug |
| PAAD | KRAS | 0.74 | 0.3554 | Cobimetinib, Binimetinib, Trametinib |
| SKCM | NRAS | 0.76 | 0.0234 | Binimetinib |
| THCA | BRAF | 0.90 | 0.4636 | Vemurafenib, Dabrafenib, Trametinib |

**Table S8. Reference WSIs used for color normalization, Related to Methods.**

| **cancer type** | **reference WSI** |
|---|---|
| Stomach adenocarcinoma | TCGA-3M-AB46-01Z-00-DX1.svs |
| Pancreatic adenocarcinoma | TCGA-2A-A8VL-01Z-00-DX1.svs |
| Prostate adenocarcinoma | TCGA-2J-AAB6-01Z-00-DX1.svs |
